# SLC26A11 is an atypical solute carrier with dual transport-channel function mediating lysosomal sulfate transport

**DOI:** 10.1101/2025.08.17.670773

**Authors:** Benedikt T. Kuhn, Peter Kovermann, Bassam G. Haddad, Tim Rasmussen, Tamsanga Hove, Stephanie Bungert-Plümke, Bettina Böttcher, Jan-Philipp Machtens, Christoph Fahlke, Eric R. Geertsma

## Abstract

Membrane transporters and channels are generally assumed to be based on distinct structural and functional principles. SLC26A11, a solute carrier with high expression levels in the brain, has been proposed to function as either an anion transporter or a channel. Here, we resolve this apparent discrepancy by demonstrating that SLC26A11 is a dual-function protein capable of operating as both a sulfate transporter and a chloride channel. By resolving its structure and combining biochemical studies and molecular dynamics simulations, we show that SLC26A11 exhibits all the hallmarks of a secondary transporter. The mechanistic basis for its selective ion transport identifies the protein as the elusive lysosomal sulfate exporter. Additionally, we demonstrate that SLC26A11 exhibits an uncoupled, channel-like chloride conductance gated by proton:sulfate symport. Our finding that the chloride-conducting state arises from the transport cycle may contribute to the development of novel therapeutic strategies for treating brain edema, and the identification of its role in lysosome sulfate efflux may provide new approaches to study and treat lysosomal storage diseases.

## Introduction

SLC26A11, or KBAT for kidney and brain anion transporter, is expressed in most tissues, with the highest expression levels in the brain ^1,2^. SLC26A11 is a member of the solute carrier family 26 (SLC26) of secondary transporters of small anions ^3,4^. Initially, SLC26A11 was described as a pH-dependent transporter of sulfate, chloride, and oxalate ^1,5,6^, consistent with its close phylogenetic relationship to plant and yeast sulfate transporters ^6^. However, subsequent electrophysiological studies in HEK293 cells ^7^ and neurons ^8,9^ have suggested that SLC26A11 functions as a chloride channel. This putative chloride channel activity of SLC26A11 has been implicated in pathological neuronal swelling ^8^ and its inhibition has been proposed as a therapeutic strategy for ischemic stroke ^9^.

Membrane transporters and channels are generally assumed to be based on different functional and structural principles ^10,11^. Based on their structure, SLC26 proteins are assumed to function as transporters that operate according to an elevator-type transport mechanism ^12^. Detailed functional studies have supported this claim by demonstrating that several SLC26 family members are secondary transporters that catalyze either anion symport ^13,14^ or exchange ^15^. However, electrophysiological measurements have also demonstrated channel-like uncoupled chloride transport for other SLC26 proteins, i.e., SLC26A7 ^16–19^, and SLC26A9 ^20–23^. Thus far, it has not been possible to determine whether these uncoupled chloride fluxes are caused by a chloride-selective channel or a fast chloride uniport transport-mode ^24^.

Until now, the function of SLC26A11 has only been studied in the plasma membrane. However, proteomic ^25,26^ and transcriptomic ^27–29^ studies and confocal microscopy ^30^ have identified SLC26A11 as a lysosomal membrane protein. Lysosomes are the primary degradative compartments of the cell ^31^. Lysosomal recycling of macromolecules depends on the concerted action of lysosomal hydrolases and transporters for catabolite export to the cytosol ^32–35^. Functional defects in either lead to the accumulation of degradation products and result in lysosomal storage diseases (LSDs) ^36,37^. While lysosomal sulfatases and their causative role in LSDs are well characterized ^38^, little is known about the efflux of the resulting sulfate ^39^, a process critical for preventing competitive inhibition of sulfatases ^40,41^. Although the existence of a lysosomal sulfate clearance pathway has been described ^42^, its molecular identity has remained unknown.

Here we characterize the three-dimensional structure and function of human SLC26A11. Our structures of nanobody-bound SLC26A11 reveal the most compact structure of any mammalian SLC26 transporter. SLC26A11 further exhibits all the hallmarks of a secondary transporter. By combining biochemical studies and molecular dynamics simulations, we show that SLC26A11 is tailored to function as the elusive lysosomal sulfate exporter and provide the mechanistic basis for its highly selective ion transport. We further demonstrate that SLC26A11 is also capable of an uncoupled, channel-like chloride conductance, which is gated by proton and sulfate transport. Taken together, our work identifies SLC26A11 as a protein with a dual transport-channel function. The identification of its role in lysosome sulfate efflux may pave the way to new approaches to study and treat lysosomal storage diseases. Our finding that the chloride-conducting state emerges from a conformation part of the secondary transport cycle may open a path to a therapeutic strategy for the treatment of brain edema.

## Results

### SLC26A11 is a proton:sulfate/chloride exchanger

Thus far, functional characterization of SLC26A11 has been performed in cell-based assays ^1,6,7,43^. These assays exclusively study proteins inserted into the plasma membrane and permit only limited control of ion gradients. Therefore, we performed a detailed *in vitro* characterization of purified and membrane-reconstituted human SLC26A11^ΔC^, a variant lacking the C-terminal intrinsically disordered region of 23 amino acids **[Supplementary Fig. 1A-C]**. Consistent with a similar truncation mutant of murine SLC26a9 ^24^, SLC26A11^ΔC^ showed higher expression levels than full-length SLC26A11 and excellent biochemical behavior **[Supplementary Fig. 1D-G]**.

Since the related *Arabidopsis* transporter, AtSultr4;1, performs pH-dependent sulfate transport ^14^, we first studied SLC26A11^ΔC^ under similar conditions (pH_out_ 5.0; pH_in_ 7.5). We observed a time-dependent accumulation of ^35^S-sulfate in SLC26A11^ΔC^ proteoliposomes, in contrast to liposomes without protein **[Fig. 1A]**. We observed that sulfate accumulation was strongly dependent on the presence of chloride ions *in trans*. Substitution of internal gluconate for chloride increased the initial rate of sulfate transport and the final accumulation level approximately fourfold. Therefore, we included 50 mM chloride in the lumen of the proteoliposomes in subsequent measurements.

**Figure 1:**
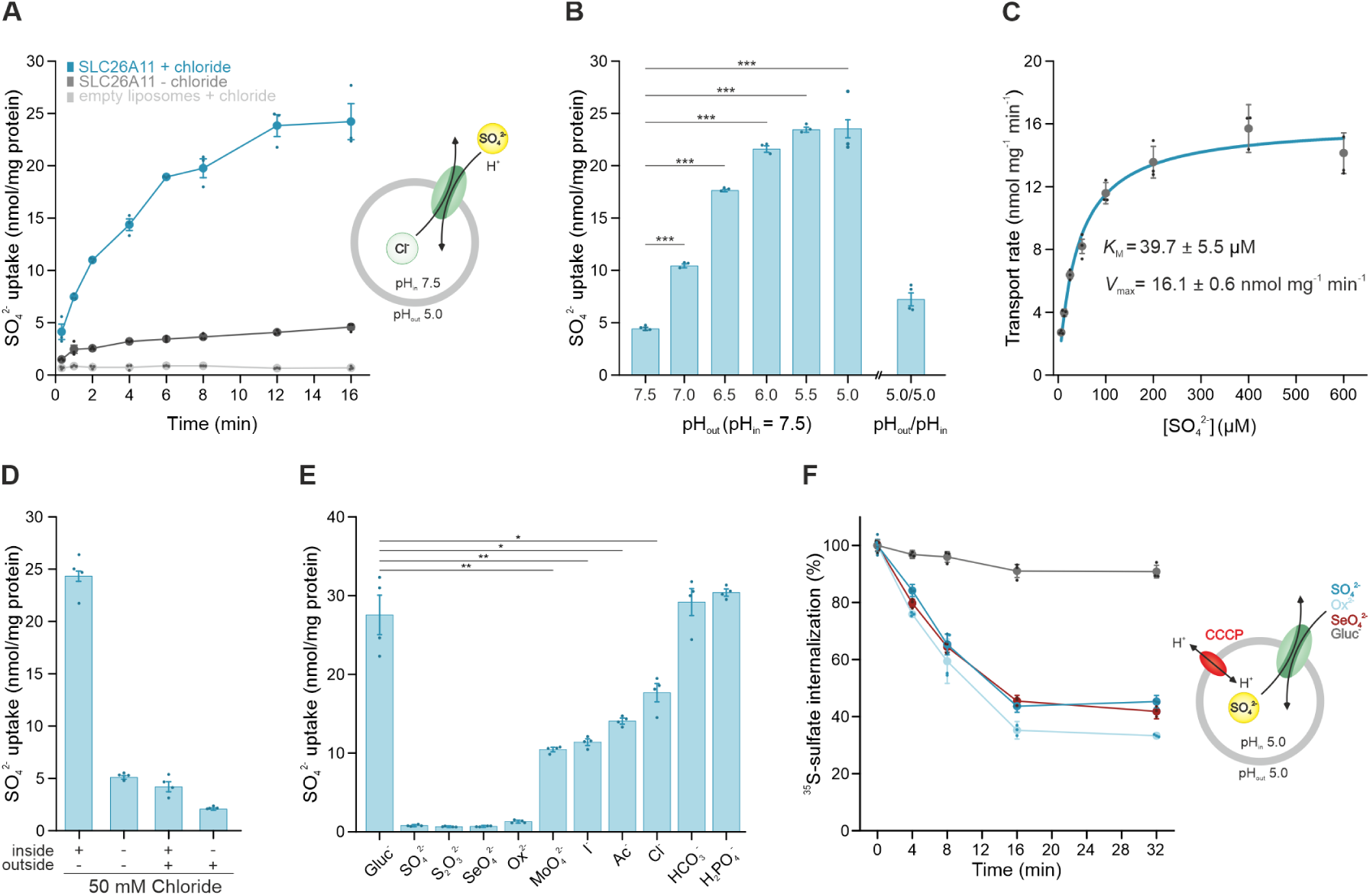
Sulfate transport of SLC26A11 is pH and chloride dependent. **(A)** Time dependent sulfate uptake into SLC26A11 proteoliposomes in presence of a pH gradient (pH 7.5 inside, pH 5.0 outside) and 50 mM internal chloride (blue) or 50 mM gluconate as chloride substitute (dark grey) and empty liposome control (light grey). **(B)** pH dependence of sulfate transport in presence of 50 mM internal chloride and with internal and external pH as indicated below the individual bars. A Two-tailed Student’s t test was performed (*** indicates p < 0.001, p values are shown in the **Source Data file**). **(C)** Sulfate transport kinetics. Data represents background-corrected initial sulfate uptake after 20 seconds in presence of increasing sulfate concentrations. K_m_ and V_max_ were derived from non-linear curve fitting using the Michaelis-Menten function in OriginPro. **(D)** Chloride dependence of sulfate transport with internal and external 50 mM chloride or 50 mM gluconate as chloride substitute as indicated below the individual bars. **(E)** Inhibition of sulfate transport in presence of different anions as indicated below individual bars. Anions were added as sodium salt at 100-fold the concentration of external sulfate. A Two-tailed Student’s t test was performed (*** p < 0.001, ** p < 0.01, * p < 0.05, p values are shown in the **Source Data file**). **(F)** Sulfate efflux from proteoliposomes upon sulfate uptake as shown in panel A. Sulfate efflux was initiated by addition of 100 nM CCCP and different anions at 100-fold external sulfate concentration as indicated. For all experiments in Fig. 1, the individual datapoints as well as the mean ± SEM (n ≥ 3) are shown. Assay buffers and precise number of replicates are detailed in **Supplementary Table 1**. Bar graphs represent background-corrected sulfate accumulation levels reached after 16 min of transport.

We further measured SLC26A11^ΔC^ sulfate transport at different pH gradients. Sulfate uptake showed a strong pH dependence, and the highest degree of sulfate accumulation was observed for pH gradients of 2.0-2.5 units and with the acidic pH outside **[Fig. 1B]**. Lower levels of sulfate accumulation were observed in the presence of a symmetrical pH of 7.5 or 5.0. Since no change in sulfate accumulation was observed in the presence or absence of an inward Na^+^-gradient, sulfate uptake does not appear to involve Na^+^ ions, in agreement with a previous study ^6^ **[Supplementary Fig. 2A]**. These results suggest that sulfate transport is obligatorily coupled to proton transport in the same direction. We determined an apparent K_M_ of 39.7 ± 5.5 µM for sulfate in the presence of a strong pH gradient (pH_out_ 5.0; pH_in_ 7.5) **[Fig. 1C]**.

Next, we systematically varied the chloride concentrations in both compartments to explain the stimulatory effect of *trans* chloride on sulfate transport **[Fig. 1D]**. While the presence of chloride in the *trans* compartment led to strong sulfate accumulation, the presence of chloride in both compartments showed similar reduced uptake levels as observed in the absence of chloride. Sulfate uptake was further reduced in the presence of chloride in the *cis* compartment only. Together, this indicates that chloride is not an allosteric modulator of SLC26A11, but rather acts as a substrate competing with sulfate for the same binding site when present in the *cis* compartment, and functions as a counter-substrate when present in the *trans* compartment.

To determine whether SLC26A11 is an obligate exchanger, we first identified anions capable of inhibiting ^35^S-sulfate uptake. We observed strong inhibition upon addition of a 100-fold excess of thiosulfate, oxalate and selenate to the *cis* compartment **[Fig. 1E]**. Molybdate, iodide, acetate and chloride inhibited transport to a lesser extent, whereas phosphate and bicarbonate had no effect on SLC26A11^ΔC^ activity. Next, we loaded proteoliposomes with ^35^S-sulfate until steady state was reached before dissipating the proton gradient. In the absence of external substrate, the proteoliposomes retained ∼90% of the sulfate over the next 32 minutes **[Fig. 1F]**. However, upon addition of a 100-fold excess of unlabeled sulfate, oxalate, or selenate, we did observe efflux of most of the ^35^S-sulfate from the proteoliposomes. The limited degree of ^35^S-sulfate efflux in the absence of a counter-substrate suggests that the energetic barrier for a conformational transition in the absence of substrate is very high. Thus, SLC26A11 appears to function most efficiently as an exchanger.

Combined these results suggest that SLC26A11 can transport protons and sulfate ions in one direction and chloride ions in the opposite direction. To approximate the stoichiometry of transported ions, we measured (*cis*) proton- and (*trans*) chloride-driven sulfate accumulation in the presence of different membrane potentials **[Supplementary Fig. 2B]**. We observed only modest changes when generating a membrane potential of 0 and +60 mV, but at -60 mV sulfate accumulation was reduced by approximately 30%. For electrogenic transport, a more pronounced reduction at -60 mV and a concomitant increase in sulfate accumulation at +60 mV would be expected, as shown for the proton:fumarate symporter SLC26Dg ^13^. These results therefore support an electroneutral transport mode for SLC26A11. In the most parsimonious model, SLC26A11 is an exchanger that couples the symport of one proton and one sulfate ion (net charge of -1) to the antiport of one chloride ion (net charge of -1).

### Overall structure of human SLC26A11

We determined structures of SLC26A11^ΔC^ in MSP1-E3D1 nanodiscs supplemented with phosphatidylcholine to gain insight into the mechanistic basis of sulfate transport coupled to the *cis* proton- and *trans* chloride-gradients. As fiducial markers for cryo-electron microscopy, we selected two nanobodies, Nb4 and Nb11, from alpaca immune libraries based on their opposite effects on the transporter’s thermal stability **[Supplementary Fig. 3]**, resulting in two highly similar structures (RMSD of 0.56 Å) with an overall resolution of 2.8 Å (Nb4) and 3.2 Å (Nb11) **[Fig. 2A, Supplementary Fig. 4-7]**. Nb4 and Nb11 each bind SLC26A11^ΔC^ on the side of the membrane domain that faces the lysosomal lumen or the extracellular environment, but they bind in different locations.

**Figure. 2:**
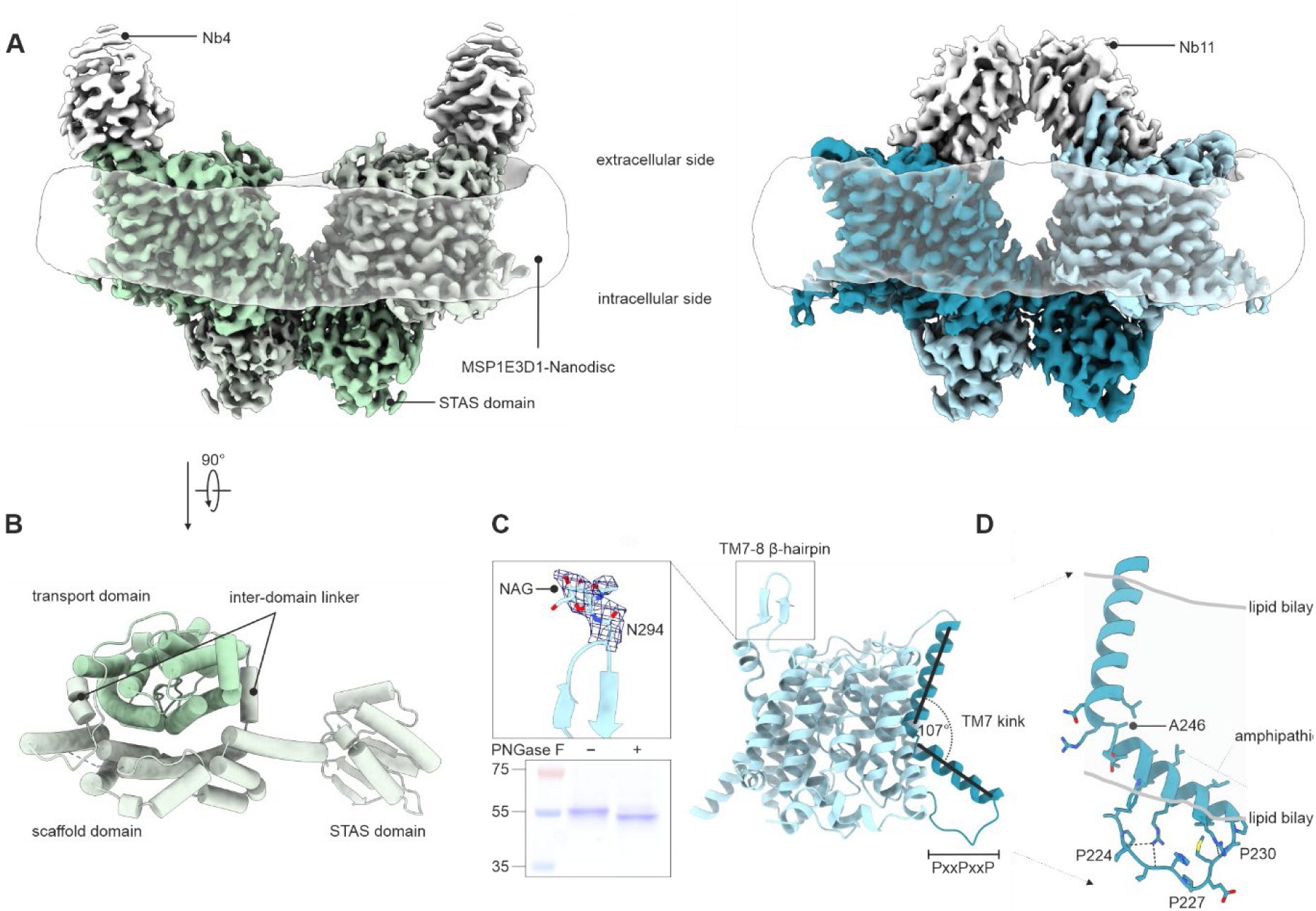
Structures of human SLC26A11^ΔC^ reveal three unique features. **(A)** Cryo-EM reconstruction of the dimeric SLC26A11^ΔC^-Nb11 (blue) and SLC26A11^ΔC^-Nb4 (green) complexes embedded in an MSP1-E3D1 nanodisc. Nanobody densities are shown in grey and the MSP1-E3D1 density, indicative of the position of the lipid bilayer, is shown as transparent belt. **(B)** View on the extracellular side of SLC26A11 ^ΔC^ showing the transport (green) and scaffold domains (light green) connected by two interdomain linkers and the intracellular STAS domain. **(C)** Alternative N-glycosylation site in SLC26A11 in the β-hairpin loop TM7-8. The electron density corresponding to the NAG group attached to Asn-294 is shown as blue mesh. Inset: PNGase F treatment increases the electrophoretic mobility of SLC26A11^ΔC^. **(D)** Helix kinking of TM7 at Ala-246 results in an extended conformation of loop TM6-7 containing the PxxPxxP SH3 binding motif. The position of the lipid bilayer, derived from the nanodisc density, is indicated as grey lines.

Both structures of nanobody-bound SLC26A11^ΔC^ show an SLC26A11 homodimer with cytoplasmic STAS domains swapped between the protomers and two nanobodies binding the membrane domain on the extracellular side. The SLC26A11 membrane domain is composed of 14 transmembrane segments (TMs) arranged in two inverted repeats of seven TMs, a fold shared by the SLC4, SLC23 and SLC26 families ^44^. The membrane domain can be further subdivided into a compact transport domain flanked on one side by an elongated scaffold domain **[Fig. 2B]**. On either side of the membrane, these subdomains are connected by α-helical interdomain-linkers that coordinate the relative reorientation of the transport domain that underlies solute transport ^45^. The structure of SLC26A11, while conforming to the overall design ^12,15,24,46–52^, deviates markedly from other mammalian SLC26 proteins in each of the three subdomains **[Supplementary Fig. 8]**.

In the transport domain, all other human isoforms carry one or more glycosylation sites in the elongated, extracellular loop bridging TM3-4. This loop is kept compact in SLC26A11 **[Supplementary Fig. 8A]**. The SLC26A11^ΔC^-Nb11 structure instead shows a β-hairpin extension in loop TM7-8, immediately adjacent to TM3-4. This hairpin is in close contact with Nb11. At its tip, this loop carries an additional density at the known glycosylation site Asn-294 ^53^, where we modeled an N-acetylglucosamine (NAG) moiety **[Fig. 2C]**. We confirmed the presence of N-glycosylation in our SLC26A11 sample by PNGase F treatment **[Fig. 2C]**. The TM7-8 loop is not resolved in the SLC26A11^ΔC^-Nb4 structure. Given that Nb4 does not interact with the hairpin, this suggests that it is highly mobile in the absence of a binding partner.

The second structural deviation in SLC26A11 involves TM7 in the scaffold domain **[Fig. 2D]**. Compared to other human SLC26 proteins, TM7 is comparably long and is kinked within the membrane so that the intracellular half is at an angle of 107° to the extracellular half **[Supplementary Fig. 8A]**. The break in the helix occurs at Ala-246 and appears to be stabilized by positioning the intracellular amphipathic half of TM7 at the membrane interface in the lipid bilayer. As a consequence of this arrangement, the intracellular loop TM6-7 is extended, presenting a potential protein-protein interaction motif, a class VIII SH3 domain binding sequence ^54^.

Finally, SLC26A11 contains the most compact STAS domain of all human SLC26 transporters **[Supplementary Fig. 8A].** This is due to the absence of an internal intrinsically disordered sequence, the so-called intervening sequence, and a comparatively short disordered C-terminus **[Supplementary Fig. 1]**. In contrast to other human SLC26 proteins, the interface between adjacent STAS domains is very small **[Supplementary Fig. 9]**. Instead, the dimer interface is largely composed of interactions between the STAS domains and TM5, TM12, TM13, and the intracellular interdomain linker, which are all part of the scaffold domain.

### Molecular basis for proton coupling

The substrate binding site is located in the center of the transport domain and faces the scaffold domain. It consists of a small cavity lined by TM1 and TM8 and the short α-helical sections of TM3 and TM10 **[Fig. 3A, B]**. Both structures capture SLC26A11^ΔC^ in the same conformation with the substrate binding site accessible from the cytoplasm via a water-filled cavity, suggesting an inward-open conformation **[Fig. 3A]**. We observed an additional density in the substrate binding site, which we modelled as a putative buffer-derived chloride ion **[Fig. 3B]**. While the substrate binding site itself is stabilized by a network of hydrogen bonds, we did not observe any specific interactions between the chloride ion and the residues lining the substrate binding site. Averaged ion densities from extensive all-atom MD simulations (*vide infra*) revealed that chloride and sulfate can bind at this site, allowing for the unambiguous assignment of the observed density **[Supplementary Fig. 10, 11]**. Chloride binding thus appears to be based on electrostatic interactions involving the positive dipoles of the α-helical segments of TM3 and TM10 and the side chain of Arg-366 **[Fig. 3B]**.

**Figure 3:**
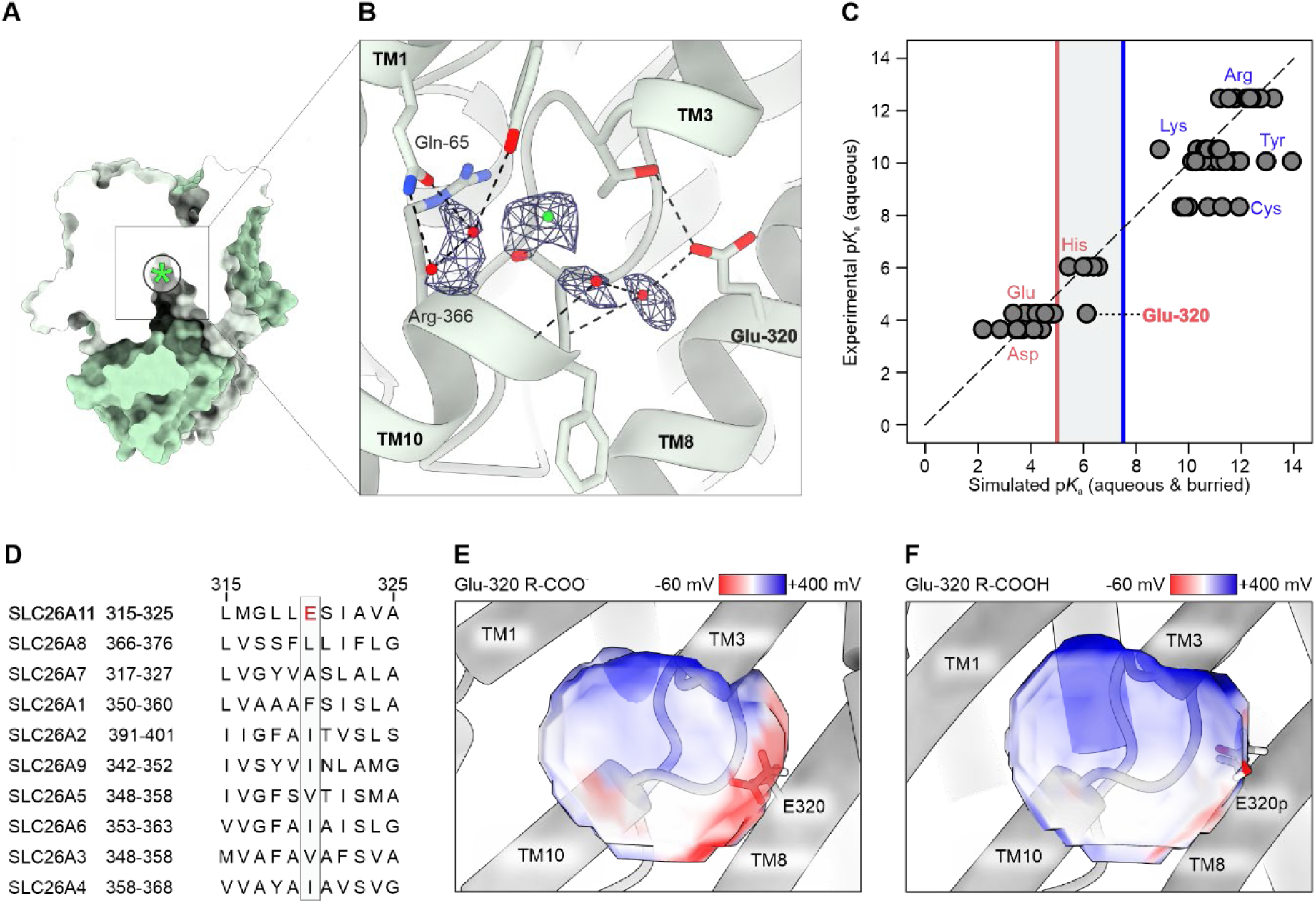
Milieu of the SLC26A11 binding site depends on a titratable residue. **(A)** Surface representation of SLC26A11^ΔC^-Nb4 clipped through the substrate binding site, indicated by a green asterisk, showing the inward-facing cavity. **(B)** Substrate binding site of SLC26A11 with bound chloride ion (green sphere), water molecules (red spheres) and corresponding electron densities shown as blue mesh. Hydrogen bonds are shown as black dotted lines. **(C)** Experimental aqueous pK_a_ values compared to simulated pK_a_ values of aqueous-exposed and buried titratable residues. The grey rectangle denotes the physiologically relevant pH range for proton-coupled transport. **(D)** Multiple sequence alignment among the human SLC26 family members, depicting a segment of TM8. The position of Glu-320 in SLC26A11 is indicated by the grey box. **(E-F)** Substrate binding site full system electrostatics in a sphere of 7-Å radius surrounding the binding site’s center-of-geometry. The substrate binding site of SLC26A11 is shown with Glu-320 in the deprotonated (E) and protonated state (F). Glu-320 is shown in stick representation.

To identify the molecular basis for proton-coupling, we performed all-atom molecular dynamics (MD) simulations of SLC26A11 with standard aqueous protonation states at pH 7 **[Supplementary Fig. 10, 11]**. For each titratable residue, we determined its average pK_a_ value over the course of the entire set of initial simulations (6.52 µs) and compared it to the corresponding standard pK_a_ value in an aqueous environment **[Fig. 3C]**. While the pK_a_ values of all histidine residues fall within the physiologically relevant pH range, these residues are located at the periphery of the protein **[Supplementary Fig. 12].** Due to the comparatively large basic shift of its pK_a_, Glu-320 also falls within the relevant pH range. Glu-320 is situated in TM8 and directly flanks the substrate binding site **[Fig. 3B]**. This analysis highlights Glu-320 as the most suitable residue for proton-coupling.

Sequence alignment of the ten human family members shows that Glu-320 is unique to SLC26A11 **[Fig. 3D]**. Although Glu-320 lines the substrate binding site, its carboxylate group appears to be too distant to contribute directly to ion coordination **[Fig. 3B]**. However, the protonation state of Glu-320 strongly affects the mean electrostatic potential in and around the substrate binding site **[Fig. 3E**, **Fig. 3F]**, consistent with a potential long-range electrostatic contribution to substrate binding.

### Glu-320 protonation controls substrate binding

The presence of a titratable residue in the substrate binding site prompted us to determine the relationship between substrate binding and pH. We first measured the binding of chloride and sulfate at pH 5.0 and pH 7.5 using differential scanning fluorimetry (DSF) and derived dissociation constants **[Fig. 4A, B, Supplementary Fig. 13]**. Chloride binding to SLC26A11^ΔC^ was independent of pH with apparent K_D_ values of 6.0 ± 1.4 and 5.3 ± 0.7 mM at pH 5.0 and pH 7.5, respectively. For sulfate we observed strong binding at pH 5.0 with an apparent K_D_ of 57 ± 11 µM. However, at pH 7.5, sulfate binding was less favorable with an apparent K_D_ of 2.9 ± 0.4 mM. Thus, while chloride binding remained constant, the affinity for sulfate changed by almost two orders of magnitude as a function of pH.

**Figure 4:**
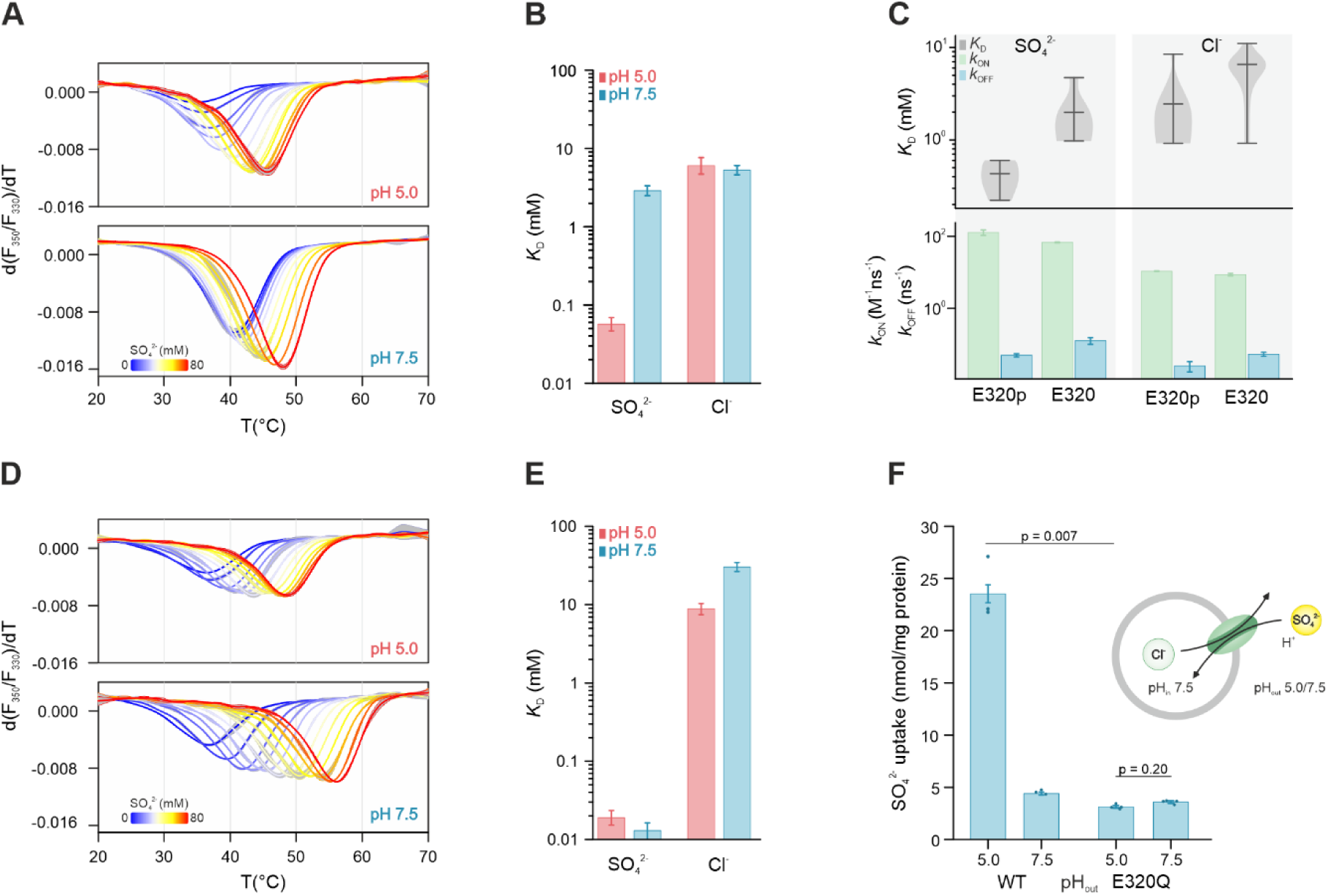
Sulfate binding to SLC26A11 is selectively controlled by the protonation state of Glu-320. **(A)** Thermal unfolding of SLC26A11^ΔC^ at pH 5.0 (top) and pH 7.5 (bottom) in the presence of increasing sulfate concentrations (blue = 0 mM sulfate, red = 80 mM sulfate). Shown are mean and standard deviation (n = 3) of the first derivative of F_350_/F_330_. The melting temperature T_m_ is reported by the local minimum of d(F_350_/F_330_)/dT. **(B)** pH-dependence of the substrate binding affinity (*K*_D_) of SLC26A11^ΔC^ and standard deviation derived from non-linear curve fitting using OriginPro of T_m_/T_m_-apo derived from panel A and **[Supplementary** Fig. 13]. **(C)** Binding affinity (*K*_D_) and binding rates (*k*_on_, *k*_off_), for sulfate and chloride, derived from all-atom equilibrium molecular dynamics simulations under different protonation states of Glu-320. E320p denotes the protonated species. Distributions of values come from six independent dimer simulations for each condition, with each subunit considered independent, resulting in 12 independent replicates per calculation. **(D)** Thermal unfolding of SLC26A11^ΔC^(E320Q) as in panel A. **(E)** pH-dependence of the *K*_D_ and standard deviation of SLC26A11^ΔC^(E320Q) as shown in panel B. **(F)** Sulfate transport of SLC26A11^ΔC^ as shown in Figure 1B and SLC26A11^ΔC^(E320Q) in presence or absence of a proton gradient as indicated. Shown are background corrected mean, SEM and individual data points (n = 3) after 16 minutes of transport. A Two-tailed Student’s t test was performed.

Next, we used DSF to analyze the pH-dependent binding of other mono- and divalent anions that act as SLC26A11 substrates **[Fig. 1E, F]**. As with chloride, we did not observe a strong pH-dependent change in binding affinity for iodide and acetate, which are completely and predominantly mono-anionic, respectively, under these conditions **[Supplementary Fig. 14]**. In contrast, the divalent anions oxalate, thiosulfate, selenate and molybdate showed a strong decrease in binding affinity, upon shifting from pH 5.0 to pH 7.5, similar to that observed for sulfate. Taken together, this suggests that the charge of the anion is the major determinant of pH modulation of binding affinity.

Given the impact of the protonation state of Glu-320 on the surface potential of the substrate binding site **[Fig. 3E, F**], we determined the sulfate and chloride binding constants for protonated and deprotonated Glu-320 using equilibrium MD simulations. Over a timespan of ∼15 µs, we captured hundreds of spontaneous binding and unbinding events, which we used to evaluate the kinetic binding constants **[Fig. 4C]**. Protonation of Glu-320, which occurs under acidic conditions, results in a slight increase in *in silico* association rates (*k*_ON_) for chloride and sulfate, and a slight decrease in the dissociation rates (*k*_OFF_) of both ions. However, both changes are more pronounced for sulfate, resulting in a significant change in binding affinity only for this ion. The dissociation constant for sulfate is reduced by almost an order of magnitude upon protonation of Glu-320. These results are in good quantitative agreement with the biochemical experiments, which also show an increased affinity for sulfate, but not for chloride, upon protonation, and demonstrate that the protonation state of Glu-320 controls the substrate-specific binding affinity.

As a mimic of the protonated state, we introduced the E320Q mutant, designated SLC26A11^ΔC^(E320Q). The chloride dissociation constant of SLC26A11^ΔC^(E320Q), as determined by DSF, was comparable to that of the wild type protein at pH 5.0 (8.8 ± 1.4 mM) and was increased six-fold at pH 7.5 (30 ± 4.0 mM) **[Fig. 4D, E]**. However, the most significant change was observed for sulfate. While the binding affinity remained high at pH 5.0 (K_D_ 19 ± 4.0 µM), the E320Q mutant showed a similarly high affinity at pH 7.5 (K_D_ 13 ± 3.0 µM), representing a more than 200-fold increase in binding affinity compared to the wild type protein. This further supports a critical role of the Glu-320 protonation state in the sulfate but not the chloride binding affinity, with the protonated and deprotonated species allowing sulfate binding with micromolar and millimolar affinities, respectively. The agreement between the K_D_ for sulfate at pH 5.0 **[Fig. 4B]** and the apparent K_M_ for sulfate transport **[Fig. 1C]** is consistent with sulfate being transported with a proton.

Finally, we did not observe any (*cis*) proton- and (*trans*) chloride-driven sulfate accumulation in SLC26A11^ΔC^(E320Q) proteoliposomes **[Fig. 4F]**. This is consistent with the very high, pH-independent sulfate binding affinity of the mutant and the slight decrease in chloride binding affinity at high pH, which together are expected to result in a very low efficiency of sulfate-for-chloride exchange in the liposomal lumen.

### SLC26A11 is a chloride channel

Since previous electrophysiological studies have suggested that SLC26A11 can function as an anion channel ^7–9^, we next studied the electrophysiological properties of SLC26A11 via whole-cell patch clamp. For these experiments we used *Spodoptera frugiperda* (Sf9) insect cells: whereas SLC26A11 exclusively localizes to lysosomes in various mammalian cell lines ^30^, a fraction of the SLC26A11 protein localizes to the plasma membrane in Sf9 cells **[Supplementary Fig. 15].**

At neutral pH and in the absence of sulfate, we did not observe any SLC26A11-mediated current above background **[Supplementary Fig. 16]**. However, in the presence of 0.5 mM sulfate in the internal solution, we detected small SLC26A11 currents (-114 ± 18 pA at -120 mV) at symmetrical chloride concentrations **[Fig. 5A-B]**. These currents exceeded the background currents in non-transduced cells (-56 ± 14 pA at -120 mV) **[Fig. 5C]**, suggesting that the observed current is conducted by SLC26A11.

**Figure 5:**
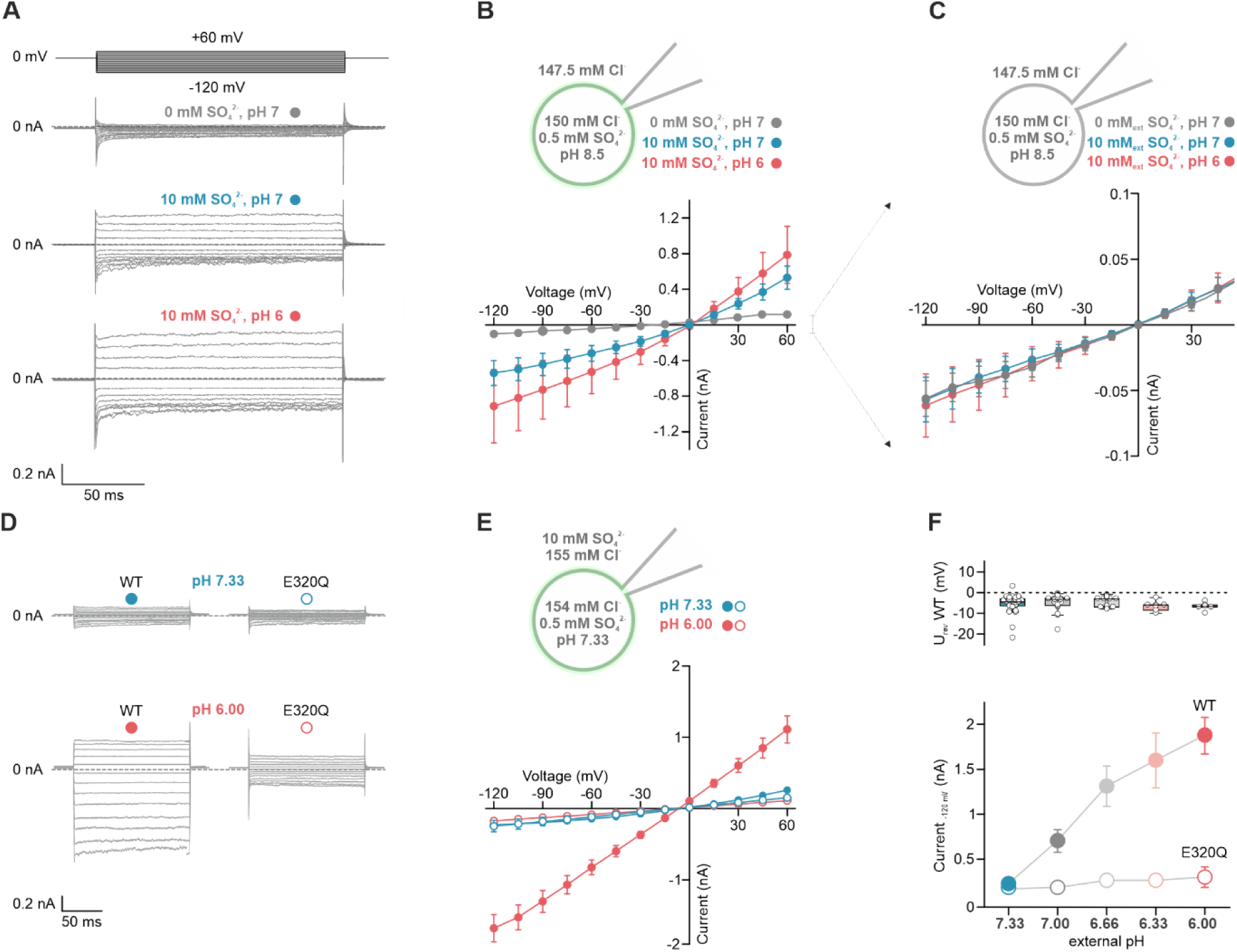
SLC26A11 exhibits pH- and sulfate-dependent intrinsic chloride channel activity. **(A)** Representative current recordings from Sf9 cells expressing SLC26A11(WT)-eGFP with buffer conditions as shown in panel B. **(B)** Current-voltage relationships (means ± standard errors; n = 13/11/7) from transfected Sf9 cells in three different bath solutions as indicated (grey: 0 mM sulfate, pH 7; blue: 10 mM sulfate, pH 7; red: 10 mM sulfate, pH 6.0). **(C)** Current-voltage relationships (means ± standard errors; n = 9/10) from untransfected Sf9 cells in the same solutions as indicated in panel B. **(D, E)** Representative whole-cell current traces **(D)** and current-voltage relationships (mean ± standard errors; WT: n = 29/6, E320Q: n = 25/11) **(E)** for SLC26A11(WT) and SLC26A11(E320Q) in absence or presence of a pH gradient (blue: symmetrical pH 7.33; red: pH_out_ 6.00, pH_in_ 7.33). **(F)** *upper panel*: Reversal potentials (U_rev_) for SLC26A11(WT) at different external pH values presented as whisker-box plots. Boxes indicate upper and lower 75/25% quartiles with medians. Whiskers span 95% confidence limits. Data points are derived from experiments shown in panel D and E and Supplementary Fig. 19. *lower panel*: Current amplitudes of SLC26A11(WT) and SLC26A11(E320Q) at -120 mV at experimental conditions as shown in panel E at various external pH conditions (pH 7.33/7.00/6.66/6.33/6.00; WT: n =29/20/12/7/6, E320Q: n = 25/15/12/12/11).

SLC26A11-specific current amplitudes further increased in the presence of 10 mM external sulfate (-463 ± 140 pA at -120 mV) and upon subsequent acidification of the bath solution from pH 7.0 to 6.0 (-833 ± 374 pA at -120 mV) **[Fig. 5A-C]**. While the latter conditions are in principle compatible with sulfate binding and transport into cells, currents of similar magnitude were also detected under conditions with opposite ion distributions compatible with sulfate binding on the cytoplasmic side and subsequent export **[Supplementary Fig. 17]**. Under conditions with high internal sulfate concentrations, currents further increased with a stronger outward pH gradient (pH_in_ 6.0; pH_out_ 8.5). Thus, conditions that favor proton:sulfate transport enhance SLC26A11 currents. Under all conditions the current reversal potential remained close to the Nernst equilibrium potential for chloride, indicating that the observed current is chloride-selective. This is further supported by the altered reversal potential observed upon depletion or substitution of external chloride **[Supplementary Fig. 18]**.

Given the impact of Glu-320 protonation on substrate selectivity and transport, we examined the consequences of the E320Q mutation on SLC26A11-specific whole-cell currents. SLC26A11(E320Q) showed similar expression levels and intracellular distribution as the wild type protein **[Supplementary Fig. 15]**. However, whereas SLC26A11(WT)-specific currents were activated by external acidification at high external sulfate, SLC26A11(E320Q) anion currents instead remained low but exceeded background currents **[Fig. 5D-F, Supplementary Fig. 19]**. The lack of pH-dependence observed for SLC26A11(E320Q) chloride currents suggests that activation of the chloride channel requires the protein to cycle through different Glu-320 protonation states, as previously observed for the electroneutral proton:sulfate/chloride exchange mode of transport **[Fig. 4F]**.

Taken together, our findings indicate that SLC26A11 exhibits two distinct transport functions: electroneutral proton:sulfate/chloride exchange and chloride channel activity. In conjunction with the sulfate dependency of chloride channel activation, the distinct pH dependence of the currents recorded with internal **[Supplementary Fig. 17]** or external sulfate **[Fig. 5A-C]** demonstrate that SLC26A11 chloride channels are gated by SLC26A11 sulfate transport.

## Discussion

The function of SLC26A11 has remained disputed. Cell-based assays have suggested that the protein is either a pH-dependent secondary transporter for sulfate, chloride, and oxalate ^1,5,6^, or a channel for chloride ^7–9^. Here, by combining functional studies performed under well-defined conditions with single-particle cryo-EM and molecular dynamics simulations, we demonstrate that human SLC26A11 can function as both a secondary-active transporter and an anion channel. This dual-function behavior resolves the remaining controversy over the function of SLC26A11 and provides the first insights into the mechanism of passive transport in this family.

Our *in vitro* transport studies using membrane-reconstituted protein indicate that SLC26A11 operates as an electroneutral proton:sulfate/chloride exchanger **[Fig.1]**. SLC26A11 thus closely resembles the function of the rat lysosomal sulfate transporter reported three decades ago ^42^. Together with its lysosomal localization ^26,30^ and its upregulation during lysosomal biogenesis ^27^, our functional characterization suggests that SLC26A11 represents the elusive lysosomal transporter for sulfate. Its use of the proton gradient as driving force to export sulfate from the lysosome is consistent with other lysosomal transporters ^37^, e.g., SLC17A5 ^55^, SLC36A1 ^56^, SLC46A3 ^57^, SLC63A1 ^58^, and SLC66A4 ^59^. In agreement with its role as a housekeeping protein, SLC26A11 shows a broad tissue distribution in the human body ^6^.

To function efficiently as a sulfate exporter, SLC26A11 must preferentially bind sulfate over chloride. This is not trivial, as the lysosomal chloride concentrations are high (80 – 120 mM) ^60,61^ and exceed the lysosomal sulfate concentrations ^40,41^ by orders of magnitude. Our results suggest that the molecular basis for this preferential binding of sulfate in the lysosomal lumen is Glu-320. Glu-320 is the only amino acid in the membrane domain of SLC26A11 that is predicted to have a pK_a_ in the physiologically relevant range **[Fig. 3C]**. Within the mammalian SLC26 family, SLC26A11 is the sole member with an acidic residue at this position **[Fig. 3D]**. Glu-320 flanks the substrate binding site **[Fig. 2]**, allowing it to contribute directly to the electrochemical milieu of the binding site **[Fig. 3E-F]**. Protonation of Glu-320 increases the binding affinity for sulfate from an apparent K_D_ of 2.9 mM to 57 µM, whereas chloride binding is essentially unaffected and remains at a comparably low affinity (apparent K_D_ values of 6.0 mM and 5.3 mM at pH 7.5 and 5.0, respectively) **[Fig. 4]**. Thus, Glu-320 acts as a signal integration node that determines whether sulfate or chloride will be favored for transport **[Fig. 4, Supplementary Fig. 13]**.

Taken together, the following molecular mechanism for transport over the lysosomal membrane emerges **[Fig. 6]**. At a lysosomal pH of approximately 4.6 ^33^, Glu-320 is predominantly protonated **[Fig. 3C]**. This selectively increases the binding affinity for sulfate **[Fig. 4B]** and allows the formation of sulfate-bound SLC26A11 **[Supplementary Fig. 20A]** despite the high lysosomal chloride concentration. While sulfate binding puts the protonated carrier in a translocation-competent state that reorients efficiently **[Fig. 1A, 1F]**, it is unclear whether the chloride-bound protonated state is mobile. Efficient reorientation of this state seems unlikely because the resulting proton leak would dissipate both the pH gradient and the membrane potential, thereby disrupting lysosomal lumen homeostasis. After reorientation of the sulfate-bound protonated binding site toward the cytoplasm, the neutral pH leads to deprotonation of Glu-320. This reduces the sulfate binding affinity and promotes sulfate unbinding **[Fig. 4B]**. Sulfate rebinding is less favorable than chloride binding at this stage **[Supplementary Fig. 20B]**. At low cytoplasmic chloride concentrations, a significant fraction of the carrier remains in the apo state. Although deprotonated *apo* SLC26A11 appears to be translocation competent, its reorientation seems much slower than that of the chloride-bound state **[Fig. 1A]**. Finally, upon subsequent arrival in the lysosomal lumen, chloride is released and Glu-320 is rapidly protonated, which may lock the carrier until a new sulfate ion is bound.

**Figure 6:**
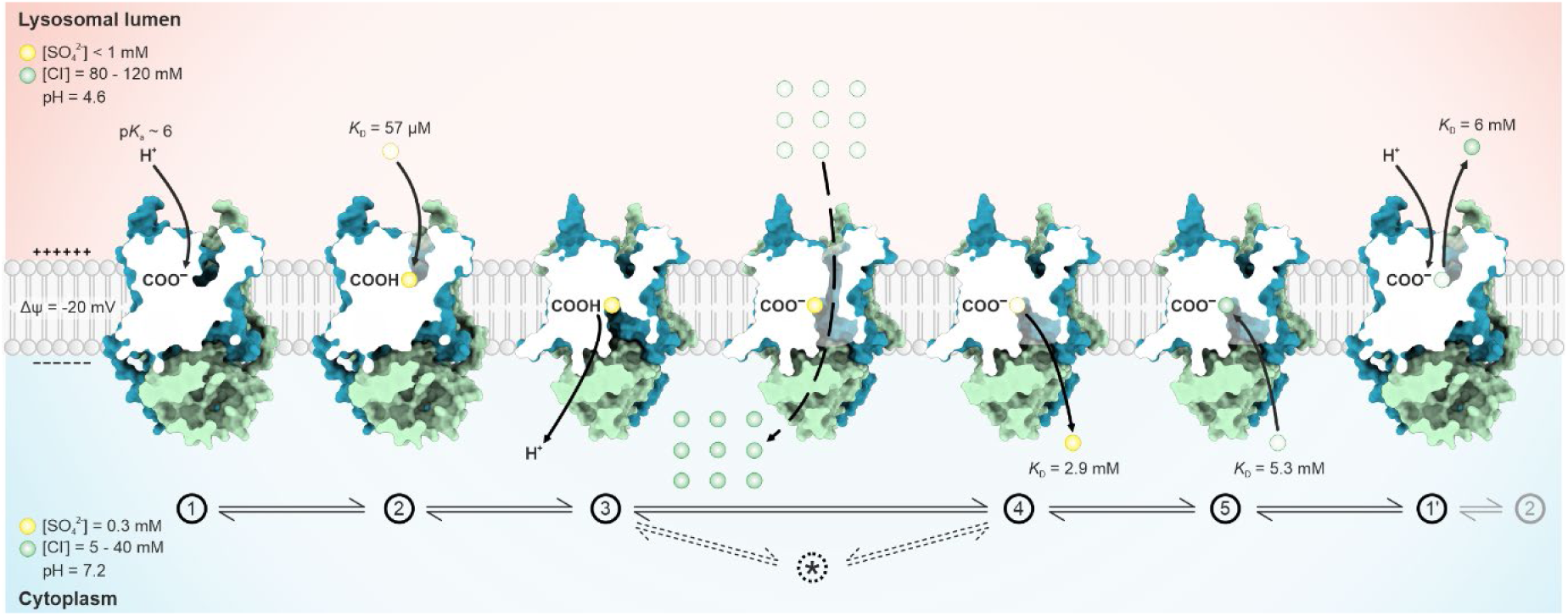
Model of the SLC26A11 dual-function mechanism. The SLC26A11 transport cycle can be broken down in five steps. (Step 1) Protonation of Glu-320 in the lysosomal lumen allows (Step 2) selective binding of sulfate despite an excess of competing chloride. Upon reorientation of the binding site to the cytoplasm, (Step 3) Glu-320 is deprotonated and (Step 4) sulfate unbinds. (Step 5) Subsequent binding of chloride accelerates the return of the substrate binding site to the lumen. In the lysosomal lumen, (Step 1’) chloride unbinds and Glu-320 is rapidly protonated, thereby completing the transport cycle. The formation of the channel-like, chloride-conducting state (Step *) requires the binding of a proton (Step 1) and sulfate (Step 2) first. It may be reached following (Step 3) the subsequent release of the proton, thereby explaining the strong reduction in conductance of the E320Q mutant, which permanently mimics the protonated state of Glu-320. Values for lysosomal and cytoplasmic sulfate and chloride concentrations, pH, and membrane potential are based on estimates ^40,41^ and published values ^33,60–62^. Representations of SLC26A11 with the substrate binding site open to the lysosomal lumen or with a channel-like configuration are toy models.

In addition to proton:sulfate/chloride exchange, heterologous expression and whole-cell patch clamp assays assigned passive chloride currents to SLC26A11 **[Fig. 5]**. In principle, passive currents can be conducted by ion channels, but also by transporters, i.e., by chloride uniporters. Since SLC26A11 functions as a proton:sulfate/chloride exchanger, a straightforward explanation for conductive chloride transport is a variation of the transport cycle resulting in chloride uniport. For example, after chloride translocation, the transporter could return to its initial conformation in the *apo* state instead of the proton- and sulfate-bound state. However, the experimentally observed enhancement of chloride currents by conditions favoring sulfate transport **[Fig. 5; Supplementary Fig. 17]** is incompatible with this uniport mode of chloride transport. The co-existence of secondary active proton:sulfate/chloride exchange and passive chloride currents activated by sulfate transport leaves little alternatives to a channel-like conducting state in SLC26A11.

Although membrane channels and transporters are traditionally considered to operate based on fundamentally different structural and mechanistic principles ^10,11^, a few other transporter families, i.e., SLC1 ^63–66^, SLC6 ^67,68^, and ClC-type transporters ^69–71^, have been shown to exhibit a similar dual-function phenotype. Among these, members of the SLC1 family of excitatory amino acid transporters have been studied in the most detail. Like SLC26 proteins, SLC1 proteins operate according to an elevator-type transport mechanism, wherein a mobile transport domain traverses the membrane relative to an immobile scaffold domain ^72,73^. In SLC1 proteins, the anion-selective conduction pore is transiently formed at the interface between the transport and scaffold domain ^74,75^. We presume that a similar mechanism underlies channel formation in SLC26A11 **[Fig. 6]**. In SLC26A11, conductive states are activated by conditions compatible with sulfate binding and transport **[Fig. 5; Supplementary Fig. 17]**. The impaired chloride currents observed for the SLC26A11(E320Q) mutant, which permanently mimics the protonated state and consequently binds sulfate with high affinity regardless of the pH, suggest that the conductive state is formed after deprotonation but before sulfate release **[Fig. 6]**.

While the relevance of sulfate export from the lysosome is apparent, the necessity of the SLC26A11 chloride channel is not. In lysosomes, the Nernst potential for chloride is likely close to the membrane potential ^33^. Thus, chloride fluxes over the SLC26A11 channel are expected to be limited. However, if the luminal chloride concentration exceeds the range that can be maintained by the membrane potential, the chloride conductance of SLC26A11 may serve to decrease its value. Given the importance of chloride in accelerating the transport mode, this feedback would ensure high transport rates.

Compared to its lysosomal counterpart, SLC26A11 residing in the plasma membrane is less likely to reach the chloride-conductive state under physiological conditions. The neutral pH and sub-millimolar sulfate concentrations on either side of the membrane ^62,76^ will suppress the formation of the protonated and sulfate-bound state that precedes the chloride-conductive conformation. However, consistent with its proposed role in pathological neuronal swelling in brain edema ^8^, high SLC26A11 chloride currents are likely to occur during brain trauma and ischemia due to the concomitant tissue acidification with external and internal pH values falling below 6.5 ^77,78^.

The subcellular localization of SLC26A11 is key to assess its impact on cellular physiology, but how SLC26A11 trafficking is orchestrated remains unclear. SLC26A11 lacks typical tyrosine or dileucine based lysosomal targeting sequences ^79^. Its appearance at the cell surface suggests that it is first trafficked from the *trans*-Golgi network to the plasma membrane and subsequently delivered to the lysosome following internalization ^80^. However, the efficiency of this process appears to vary between cell types, as SLC26A11 is not consistently observed in the plasma membrane ^30^. Glycosylation does not affect the plasma membrane localization of SLC26A11 ^53^, in contrast to other lysosomal membrane proteins ^81^. Alternatively, association with other membrane proteins may modulate transport between compartments ^82^. While heterodimerization with other SLC26 proteins has been ruled out ^30^, SLC26A11 has been shown to co-localize with the vacuolar-type H^+^-ATPase (V-ATPase) ^1^ and modulate its activity ^7^. Since the SLC26A11-STAS domain is remarkably compact and lacks the disordered sequences typically used for protein-protein interactions in SLC26 proteins ^4^, the unique SH3 domain binding motif **[Fig. 2D]** between TM6 and the elongated kinked TM7 may contribute to such an association with a lysosomal membrane protein.

In conclusion, our structural and functional characterization demonstrate that human SLC26A11 is a dual-function protein that operates as both a coupled secondary transporter for small anions and a chloride channel. The physiological relevance of these two modes depends on the subcellular localization of the protein. SLC26A11 in lysosomes is the elusive sulfate export system and its chloride conductance contributes to lysosomal chloride homeostasis thereby ensuring high proton:sulfate/chloride exchange rates. Under physiological conditions, the activity of plasmalemmal SLC26A11 is limited. Only upon tissue acidification caused by, e.g., ischemia, SLC26A11 is activated as a chloride channel that mediates anion influx and contributes to neuronal swelling ^8^. The proposed coupling of SLC26A11 transport and anion channel opening implies that specific transport inhibitors, such as a nanobody that binds to the extracellular face of SLC26A11, may contribute to a therapeutic strategy to block chloride influx through SLC26A11 and prevent cytotoxic brain edema caused by neuronal swelling.

## Material/Methods

### Molecular cloning of SLC26A11(ΔC)

The open reading frame of SLC26A11 was PCR amplified from the pDONR221_SLC26A11 plasmid (Addgene: #131996) using FX-cloning primers (https://www.fxcloning.org; for primer (5’-3’): *atatatgctcttctagtcccagctccgtgaccgccctgggacag*, rev primer (5’-3’): *atatatgctcttctagtccaagctccgtgaccgccctggga*). The column purified PCR product was cloned into pINIT_cat (Addgene: #46858) using FX cloning ^83^. The Insert was sequence-verified, subcloned into the pHXC3GS or pHXCA3GS vector and transformed into *E. coli* MC1061 for plasmid production.

### SLC26A11(ΔC, E320Q) mutagenesis

The open reading frame of SLC26A11 was PCR amplified from the pDONR221_SLC26A11 plasmid (Addgene: #131996) in 2 separate PCR reactions using the following primers: reaction 1 for primer (5’-3’): *atatatgctcttctagtcccagctccgtgaccgccctgggacag*, reaction 1 rev primer (5’-3’): *atatatgctcttcattgcagcaggcccatcaggggcaccactgc*, reaction 2 for primer (5’-3’): *atatatgctcttctcaatctatcgccgtggccaaggcctttgcc*, reaction 2 rev primer (5’-3’): *atatatgctcttctagtccaagctccgtgaccgccctggga*. PCR products of reaction 1 and 2 were column purified, combined to equal molar ratio and cloned into pINIT_cat (Addgene: #46858) using FX cloning. The insert was sequence-verified, subcloned into the pHXC3GS vector and transformed into E. coli MC1061 for plasmid production.

### SLC26A11 expression and purification

Baculovirus was prepared as described elsewhere ^84^ using the pHXC3GS or pHXCA3GS vectors harboring the SLC26A11(ΔC) open reading frame. *Trichoplusia ni* cells at a concentration of 1×10^6^ cells/mL were infected with 1% (v/v) baculovirus and cultivated in suspension for 72 h at 27 °C in ESF 921 serum-free medium. Cells were collected by centrifugation at 300 x g and 4 °C for 10 minutes and resuspended in 20 mM Hepes pH 7.25, 150 mM NaCl, 10% glycerol, 1 mM PMSF, 1 mM MgCl_2_, 2% DDM, 0.2% CHS and stirred for 1 h at 4 °C in presence of Benzonase. Cell debris was removed by centrifugation at 140000 x g and 4 °C for 30 minutes. The supernatant was submitted to batch binding with washed and pre-equilibrated Strep-Tactin® Superflow® resin (IBA) for 1 h. The sample was loaded on a gravity flow column, the column drained by gravity and the resin washed with 20 column volumes wash buffer (20 mM Hepes pH 7.25, 150 mM NaCl, 10% glycerol, 0.05% DDM, 0.005% CHS). SLC26A11 was eluted from the column by addition of 3 column volumes wash buffer supplemented with 2.5 mM D-desthiobiotin and subsequently incubated for 1 h at 4 °C in presence of 1 mg HRV-3C protease. The sample was concentrated at 2000 x g and 4 °C using an Amicon Ultra 50 kDa MWCO concentrator, centrifuged for 5 min at 25,000 x g to remove large aggregates and subsequently loaded on a Superdex 200 increase 10/300 GL size exclusion column equilibrated with buffer containing 20 mM Hepes pH 7.25, 150 mM NaCl, 0.05% DDM, 0.005% CHS. Peak fractions corresponding to dimeric and monomeric SLC26A11 were pooled and diluted to 1 mg/ml concentration in 20 mM Hepes pH 7.25, 150 mM NaCl, 10% glycerol, 0.05% DDM, 0.005% CHS, snap frozen in liquid nitrogen and stored at -80°C until further use.

SLC26A11 for alpaca immunization was prepared in buffer containing a 10-fold higher CHS concentration (20 mM Hepes pH 7.25, 150 mM NaCl, 10% glycerol, 0.05% DDM and 0.05% CHS). For nanobody selection, SLC26A11 was expressed with a C-terminal AVI-tag and enzymatically biotinylated as described elsewhere ^85^. Biotinylation efficiency was quantified using the mobility shift of biotinylated protein in SDS-PAGE upon the addition of Streptavidin and exceeded 90%.

### Nanobody selection

Nb11 was selected against SLC26A11(ΔC) following the described procedure ^86,87^. Two alpacas were immunized 4 times over a time course of 6 weeks. Four days after the final antigen injection, peripheral blood lymphocytes were isolated. The RNA was purified and converted to cDNA by reverse-transcription, the repertoire amplified by PCR and nanobody genes were cloned into the phage display compatible pDX_init phagemid ^88^ (Addgene: #110101). Two rounds of phage display were performed, binders selected based on their ELISA-signal intensity and sequence analyzed.

### MSP1E3D1 nanodisc reconstitution and cryoEM sample preparation

SLC26A11(ΔC) was expressed and purified as described above but in the absence of CHS. Glycerol was excluded from the wash and elution buffers. Affinity chromatography pure protein was mixed with DDM solubilized SoyPC lipids in a 1:50 molar ratio and incubated for 1 h at 4 °C with gentle agitation. MSP1E3D1 protein was added to a final molar ratio of 1:1:50 (SLC26A11(ΔC):MSP1E3D1:SoyPC) and the sample incubated for 5 h at 4 °C with gentle agitation. Dried Bio-Beads SM-2 were added to a final amount of 50 mg beads/1 mg DDM. The sample was incubated overnight at 4 °C with gentle agitation. Bio-Beads were removed using a gravity column and the sample collected from the eluate. The SLC26A11(ΔC)-GFP fusion protein was cleaved off for 1 h at 4 °C upon addition of 500 μg HRV-3C protease. The sample was concentrated at 3000 x g and 4 °C using an Amicon Ultra 100 kDa MWCO concentrator. The concentrated sample was centrifuged for 10 min at 13,000 x g to remove large aggregates and subsequently loaded on a Superose 6 increase 10/300 GL size exclusion column equilibrated with 20 mM Hepes pH 7.25, 150 mM NaCl. The main peak fraction was applied to a second size exclusion run using the same column and buffer. The main peak fraction was concentrated at 7000 x g and 4 °C using an Amicon Ultra 0.5 mL 50 kDa MWCO concentrator. Purified Nb11 was added to the size exclusion chromatography pure SLC26A11(ΔC) nanodiscs in a 1:1.2 molar ratio and allowed to bind for 1 h at 4 °C before freezing.

### Proteoliposome preparation

Purified SLC26A11(ΔC) was mixed with preformed, Triton X-100 destabilized large unilamellar vesicles from SoyPC and DDM solubilized BMP lipid to a ratio of 1:50:0.5 (wt/wt/wt) SLC26A11:SoyPC:BMP (LPR 50) and incubated 15 min at room temperature. Detergent was removed from the sample by stepwise addition of Bio Beads SM-2 as described previously ^89^. Bio Beads were removed from the sample using a gravity column. Proteoliposomes were collected by centrifugation for 1 h at 140000 x g and resuspended to 20 mg lipid/mL concentration in 50 mM KPi pH 7.5 using a 25-gauge needle and snap frozen and stored in liquid nitrogen until further use.

### Radioisotope transport assays

Proteoliposomes were thawed, centrifuged at 250000 x g and 15 °C for 20 minutes and resuspended in buffer containing 20 mM Hepes, 20 mM Mes pH 7.5, 2 mM Mg-gluconate and 50 mM KCl. Proteoliposomes were snap frozen in liquid nitrogen and thawed at room temperature three times before 11x extrusion using a 400 nm polycarbonate filter. Proteoliposomes were centrifuged at 250000 x g and 15 °C for 20 minutes, the supernatant discarded and the pellet resuspended to 100 mg/ml concentration in buffer containing 20 mM Hepes, 20 mM Mes pH 7.5, 2 mM Mg-gluconate and 50 mM KCl and homogenized with a 25-gauge needle. To initiate transport, proteoliposomes were diluted to 2.5 mg/ml concentration in 30 °C warm buffer containing 20 mM Hepes, 20 mM Mes pH 5, 50 mM K-gluconate, 2 mM Mg-gluconate and 50 μM K_2_SO_4_ with 2 μCi/ml activity. The transport reaction was performed in a water bath at 30 °C and stopped at regular intervals by 20-fold dilution of a 100 µl aliquot in ice-cold uptake buffer followed by rapid filtration on 0.45 µm nitrocellulose filters. Filters were washed with additional 2 ml buffer, dissolved overnight in 4 ml Ultima Gold^TM^ LSC cocktail (SigmaAldrich) and analyzed using a Tri-carb 2900TR (PerkinElmer) liquid scintillation counter.

### Cryo-EM methods

Samples were vitrified on UltrAuFoil grids (R0.6/1.0, Quantifoil, Germany) after glow discharge for 150 sec in air at a pressure of 3.0×10^-1^ Torr at medium power with a Harrick Plasma Cleaner (PDC-002). A Vitrobot IV (ThermoFisher) was used for vitrification in liquid ethane with 5 s blot time and +20 blot force. Movie data was acquired at the cryo-EM facility in Würzburg on a Krios G3 electron microscope (ThermoFisher) with different cameras and settings for the two samples: For SLC26A11 with Nb11 a Falcon III direct detector was used in counting mode, collecting 47 fractions in 75 sec. At a magnification of 75,000x a calibrated pixel size of 1.0635 Å was obtained and a total exposure of 79 electrons/Å^2^ was used. The target defocus range was between 1.0 to 1.4 µm under focus. 2553 movies were recorded and then motion corrected and dose weighted in a live session of the program package *cryoSPARC* version 4.4 ^90^ followed by patch based CTF estimation. All subsequent steps of image processing were also performed in *cryoSPARC*. The initial set of 1,325,543 particles were obtained with a blob picker and cleaned up by 2D classification to 570,899 particles. Three initial volumes were obtained by an ab-initio reconstruction. The initial volumes were used in a heterogeneous refinement as starting references. One of the resulting classes with 276,124 particles was further refined in a non-uniform refinement with C1 symmetry and used as input for a variability analysis with three principal components subdividing the data set into three clusters. One of three clusters with 48,803 particles was then analysed in another non-uniform refinement with applied C2 symmetry and reached a resolution of 3.2 Å.

For SLC26A11 with Nb4, a Falcon IVi direct detector attached to a Selectris energy filter was used. Movies were recorded as zero-loss images with a slit width of 5 eV. A magnification of 130,000x resulted in a calibrated pixel size of 0.946 Å and a total exposure of 70 electrons/Å^2^ was used with an exposure time of 6.2 s. The target defocus range was between 0.6 and 1.6 µm under focus. 11086 movies were recorded in EER format. Further image analysis was performed with *cryoSPARC*. The EER movies were converted into 40 fractions, followed by motion correction, dose weighting and averaging of the fractions. The CTF of the averaged fractions was estimated with patch ctf. Initially, 2,147,241 particles were picked with a blob picker and reduced to 115,117 particles by 2D classification keeping particles in classes with the best-defined class averages. Three initial volumes were determined in an ab-initio reconstruction. These volumes were used as starting references in a heterogeneous refinement. The classes were further refined by non-uniform refinement. One of the classes provided a 3D-map, which was projected equally in space to create templates for template picking. Template picking identified 3,616,816 particles which were reduced to 457,309 particles by 2D-classification and selection of the best 2D-classes. The selected particles were analysed by another round of heterogeneous refinement. One of the classes of the heterogeneous refinement with 210,771 particles was subjected to non-uniform refinement with imposed C2-symmetry. This resulted in a map with 2.8 Å resolution, which was then filtered according to the local resolution.

An initial atomic model of the map was obtained with the program *Modelangelo* ^91^. The model was manually optimised using *Coot* version 0.9.8.93 ^92,93^ and ChimeraX/ISOLDE ^94^ and refined with the real-space refinement tool of *Phenix* version 1.21.1 ^95^.

### Differential scanning fluorometry

SLC26A11(ΔC) was dialyzed 3 times 30 minutes at 4 °C against buffer containing 20 mM Hepes pH 7.25, 0.05% DDM to remove chloride ions from the sample. The protein was diluted to 0.1 mg/ml final concentration in buffer containing 100 mM Hepes, 100 mM MES, pH 7.5 or pH 5.0, respectively, with 80 mM of the tested anion as sodium salt and 70 mM sodium gluconate. For affinity determination of sulfate or chloride, respectively, titration of the tested anion starting from 80 mM was performed and the total anion concentration kept constant at 150 mM by addition of additional sodium gluconate. Samples were equilibrated for 30 minutes on ice, 15 minutes at room temperature and subjected to thermal melting in triplicates from 20 to 70 °C with a ramp of 1 °C/min using a Prometheus NT.48 (Nanotemper) at 100% gain and Prometheus standard capillaries. The melting curves were analyzed and Tm values calculated with the PR. StabilityAnalysis software (Nanotemper). For affinity determination, ΔTm/Tm_Apo_ was plotted against the anion concentration and a non-linear curve fit performed in OriginPro using equation 1 as described elsewhere ^96,97^:

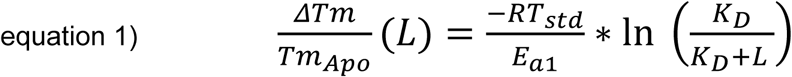

where L is the total ligand concentration; R is the universal gas constant; T_std_ is the standard temperature (298.15 K); E_a1_ the activation energy of apo state unfolding.

For affinity determination of anions other than chloride or sulfate, ΔTm/Tm_Apo_ was derived from thermal melting in presence of 80 mM substrate concentration and the dissociation constant calculated using equation 2 with E_a1_ derived from sulfate/chloride titration as described elsewhere ^97^:

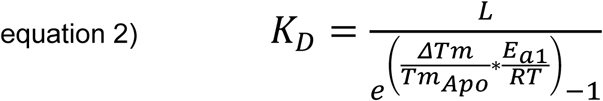

### Molecular dynamics simulations

The final coordinates of the SLC26A11 structure were used for all-atom molecular dynamics (MD) simulations. We modelled residues 26–583, neutralizing the artificial N- and C-termini using acetylation and n-methylamidation respectively using VMD ^98^. MD simulations and analysis were performed using GROMACS 2022 ^99^. An additional system was generated with glutamate 320 protonated (E320p); the protonation, remaining hydrogen atoms, and topologies were generated using *gmx pdb2gmx*. The protein was embedded in a membrane containing 1-palmitoyl-2-oleoyl-glycero-3-phosphocholine (POPC) using *gmx membed*. The initial system was solvated (TIP3P) and ionized with 200 mM NaCl, with subsequent neutralization. The protein was modelled using the CHARMM36m force-field, lipids, water, and Na^+^/Cl^-^ ions were modelled with CHARMM36. For sulfate, we took the initial parameters from Cannon *et al*. ^100^, with the addition of a non-bonded coefficient alteration (CUFIX) ^101^ to account for sulfate-protein interactions. To increase the possibility of sulfate binding we removed all chlorides, and added 15 molecules of SO_4_^2−^, resulting in a final concentration of ∼ 10 mM Na_2_SO_4_ after neutralization. We used the following minimization/equilibration protocol for the E320 (deprotonated system) with 200 mM NaCl: After embedding, the protein was minimized using the steepest descent algorithm (step 1), followed by 50 ns of simulation with position restraints on all atoms except for lipid heavy atoms in the XY plane (step 2.0). The next 50 ns step had position restraints only on the protein (step 2.1), and finally 50 ns with only protein backbone restraints (step 2.2). Equilibration was monitored via the number of protein-lipid contacts. All simulations are performed under the NPT ensemble, with v-rescale ^102^, and c-rescale ^103^ for temperature and pressure coupling respectively. Protonated systems (E320p), and systems containing Na_2_SO_4_ were constructed from the final frame of step 2.1, i.e., added proton and/or replaced 200 mM NaCl with 10 mM Na_2_SO_4_, followed by step 2.2 for each additional system – equilibration was monitored for each system separately. The final 10 ns of step 2.2 were used to initiate three replicates for the Na_2_SO_4_ containing systems, and six replicates for the NaCl systems, each replicate being run for a minimum of 700 ns totalling roughly 15 µs.

### Computational predictions of pK_a_

To approximate the pK_a_ values for titratable residues, propkatraj ^104^ was used. Due to the heuristic approach of PROPKA 3.1 ^105^, the underlying algorithm in propkatraj, pK_a_ predictions from individual conformations are subject to significant variance, and lack accuracy – propkatraj helps to ameliorate this issue by estimating the pK_a_ for many frames (1frame/ns for ∼6500 ns), resulting in distributions of pK_a_, centred around a mean-value. These average pK_a_ values were reported in Figure 3B.

### Biophysical characterization of the anion binding pocket (ABP)

To dissect the molecular interactions between chloride, sulfate, and the ABP, electrostatics of the ABP, and kinetics/thermodynamics of the anion–ABP interactions were evaluated ^106^. Full system electrostatics were calculated using g_elpot ^107^, which evaluates the electrostatic potential felt by water molecules. To this end, a sphere with a radius of 7 Å, centred at the ABP was coloured according to the electrostatic potential at the surface of the sphere [**Fig. 3E, F**].

To evaluate the kinetics of binding, we define the inner- and outer-boundary of 14 Å and 15 Å respectively, as the boundaries of a Schmitt-Trigger which filters out noise at the ABP entrance. We define anion binding according to a single-ion radial distribution function to the ABP center of geometry. The distance of 14 Å represents the first contact that the anion makes within the intracellular vestibule of SLC26A11. When an anion was within 14 Å of the ABP center of geometry, it was considered bound (1), whereas distances greater than 15 Å were considered unbound (0). When the distance is the range 14-15 Å, the bound state (1 or 0) is determined by the previous bound state. The state-trajectories are separated into *N_Bound_* groups of consecutive 1’s, where*N_Bound_* is the number of binding events.

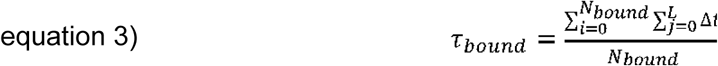

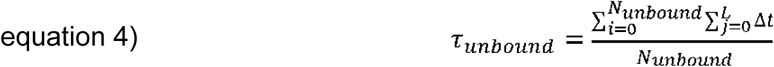

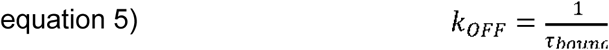

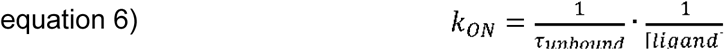

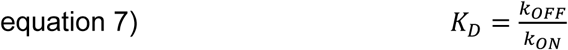

Here, *T*_*bound*_ is the average dwell-time, *L* is the length of a consecutive group of 1’s, and Δ*t* is the time-step of the state-trajectory, which is 0.1 ns for all systems. Similarly, *T*_*unbound*_, and consequently *k*_*on*_ can be calculated similarly by evaluating *N*_*unbound*_ consecutive groups of 0’s from the state-trajectory. Since *k*_*on*_ is a second order rate-constant, we normalized it to the bulk concentration of chloride (0.2 M) and sulfate (0.009 M) respectively.

### Electrophysiology

For electrophysiological characterization, wild type and mutant *h*SLC26A11 fused to GFP was expressed in *Sf*9 insect cells (*Spodoptera frugiperda*) and tested 2-5 days after infection with 1% (v/v) baculovirus. Standard whole-cell patch clamp recordings were performed with an EPC-10 USB amplifier (HEKA Elektronik, Göttingen) with pipette resistances between 3-5 MΩ as described ^108,109^. Cells were clamped to 0 mV for at least 5 s between test sweeps, and voltage pulses were applied for 150 ms from -120 mV to +60 mV in 15 mV steps. Junction potentials were corrected *a priori*. For the recordings shown in **[Fig. 5b]**, external solutions contained 128.5 mM NaCl, 7.5 mM Na-gluconate, 1 mM MgCl_2_, 5 mM CaCl_2_, 5 mM TEA-Cl, 10 mM HEPES/Tris, pH 7, or 5 mM HEPES/5 mM MES, pH 6. The pipette internal solution contained 150 mM KCl, 0.5 mM K_2_SO_4_, 2 mM MgCl_2_, 5 mM EGTA and 10 mM HEPES/KOH, pH 8.5. Anion selectivity **[Supplementary Fig. 18]** was tested by external perfusion of cells with solutions containing 0.5 mM Na_2_SO_4_ in combination with 145 mM Cl^-^ (128 mM NaCl, 0.5 mM Na_2_SO_4_, 1 mM MgCl_2_, 5 mM CaCl_2_, 5 mM TEA-Cl, 10 mM HEPES/Tris pH 8.5), 5 mM Cl^-^ (128 mM Na-gluconate, 0.5 mM Na_2_SO_4_, 1 mM Mg-gluconate, 5 mM Ca-gluconate, 5 mM TEA-Cl, 10 mM HEPES/Tris, pH 8.5) or 147 mM SCN^-^ (147 mM NaSCN, 0.5 mM Na_2_SO_4_, 1 mM MgCl_2_, 5 mM CaCl_2_, 5 mM TEA-Cl, 10 mM HEPES/Tris, pH 8.5) at pH 7, with an internal solution containing 120 mM KCl, 10 mM K_2_SO_4_, 2 mM MgCl_2_, 5 mM EGTA, 5 mM MES/5 mM HEPES/KOH, pH 6.0. For the pH dependences **[Supplementary Fig. 19]** the external solutions contained 128 mM NaCl, 10 mM Na_2_SO_4_, 1 mM MgCl_2_, 5 mM CaCl_2_, 5 mM TEA-Cl, buffered with 10 mM HEPES/Tris and MES/Tris mixtures to pH 7.33/pH 7.0/pH 6.66/pH 6.33/pH 6.0. The internal solution contained 150 mM KCl, 0.5 mM K_2_SO_4_, 2 mM MgCl_2_, 5 mM EGTA, 10 mM HEPES/Tris, pH 7.33).

## Data availability

All raw data are available upon request. The python script used in the calculation of kinetic constants from MD simulations is available at https://jugit.fz-juelich.de/computational-neurophysiology/SLC26A11-ion-binding.

## Acknowledgements

We would like to thank Barbara Borgonovo, Aliona Bogdanova, and Régis Lemaitre from the Protein Biochemistry facility of the MPI-CBG for their support. We further thank Sasa Stefanic from the Nanobody Service facility of the University of Zurich for his ongoing support. We acknowledge past and present members of FOR5046 for stimulating discussions and critical feedback. We further acknowledge Michele Marass for corrections to the manuscript. This work was funded by the Deutsche Forschungsgemeinschaft (DFG, German Research Foundation) as part of Research Unit 5046 (FOR5046; project 426950122) to E.R.G. (subproject P1), J.-P.M. (subproject P2), as well as to ChF (subproject P4), and projects 359471283, 456578072, 525040890 (EM facility Würzburg; BB). The authors gratefully acknowledge computing time on the supercomputer JURECA at Forschungszentrum Jülich under grant jara0177. All members of the Geertsma lab are acknowledged for help in all stages of the project.

## Author contributions

B.T.K., J.-P.M., Ch.F., and E.R.G. conceived the project. B.T.K. performed all protein biochemistry, carried out reconstitutions in proteoliposomes and nanodiscs, performed *in vitro* transport studies and binding assays, and selected nanobodies under the supervision of E.R.G. P.K. performed electrophysiological characterization of SLC26A11 in insect cells with the help of S.B.-G., using baculovirus provided by B.T.K, and under the supervision of Ch.F. B.G.H. performed molecular dynamics simulations, pKa predictions, electrostatic characterization, and kinetic analysis of ion binding under the supervision of J.-P.M. T.R. prepared grids, recorded, and processed cryo-EM data with the help of T.H. and under the supervision of B.B. T.R., T.H. and B.T.K. built and validated cryo-EM structures. All authors discussed and analyzed the results. B.T.K. and E.R.G. wrote the first draft of the manuscript with contributions from all authors. All authors participated in the revision of the manuscript.

