## Supplemental files for "SLC26A11 is an atypical solute carrier with dual transport-channel function mediating lysosomal sulfate transport"

Benedikt T. Kuhn *et alia*

**Supplementary Information**

- Supplementary Table 1
- Supplementary Figures 1 – 20

**Supplementary Table 1: Buffer conditions used in proteoliposome transport assays with variable component highlighted in bold.** Hepes/Mes buffer stocks were pH adjusted using potassium hydroxide if not state differently. All gluconate stocks were pH adjusted using gluconic acid.

| figure panel | internal buffers | external buffers | replicates |
|---|---|---|---|
| 1A | 20 mM Hepes, 20 mM Mes<br>50 mM potassium<br>2 mM magnesium gluconate<br>**50 mM chloride**<br>pH 7.5<br><br>20 mM Hepes, 20 mM Mes<br>50 mM potassium<br>2 mM magnesium gluconate<br>**50 mM gluconate**<br>pH 7.5 | 20 mM Hepes, 20 mM Mes<br>50 µM potassium sulfate (2 µCi/mL)<br>50 mM potassium gluconate<br>2 mM magnesium gluconate<br>pH 5.0 | n = 3 |
| 1B/4F | 20 mM Hepes, 20 mM Mes<br>50 mM potassium chloride<br>2 mM magnesium gluconate<br>**pH 5.0**<br><br>20 mM Hepes, 20 mM Mes<br>50 mM potassium chloride<br>2 mM magnesium gluconate<br>**pH 7.5** | 20 mM Hepes, 20 mM Mes<br>50 µM potassium sulfate (2 µCi/mL)<br>50 mM potassium gluconate<br>2 mM magnesium gluconate<br>**pH 5.0 to pH 7.5 (0.5 pH steps)** | n = 4 for pH5/pH5,<br>n = 3 for all remaining conditions |
| S2A | 20 mM Hepes, 20 mM Mes (KOH)<br>50 mM potassium chloride<br>2 mM magnesium gluconate<br>pH 7.5 | 20 mM Hepes, 20 mM Mes (**KOH**)<br>50 µM potassium sulfate (2 µCi/mL)<br>**50 mM potassium gluconate**<br>2 mM magnesium gluconate<br>pH 5.0<br><br>20 mM Hepes, 20 mM Mes (**NaOH**)<br>50 µM potassium sulfate (2 µCi/mL)<br>**50 mM sodium gluconate**<br>2 mM magnesium gluconate<br>pH 5.0 | n = 4 |
| 1C | 20 mM Hepes, 20 mM Mes<br>50 mM potassium chloride<br>2 mM magnesium gluconate<br>pH 7.5 | 20 mM Hepes, 20 mM Mes<br>**potassium sulfate (2 µCi/mL)**<br>**(6.25, 12.5, 25, 50 or 100 µM)**<br>50 mM potassium gluconate<br>2 mM magnesium gluconate<br>pH 5.0<br><br>20 mM Hepes, 20 mM Mes<br>**potassium sulfate (10 µCi/mL)**<br>**(200, 400 or 600 µM)**<br>50 mM potassium gluconate<br>2 mM magnesium gluconate<br>pH 5.0 | n = 3 |

| figure panel | internal buffers | external buffers | replicates |
|---|---|---|---|
| 1D | 20 mM Hepes, 20 mM Mes<br>2 mM magnesium gluconate<br>**50 mM potassium chloride**<br>pH 7.5 | 20 mM Hepes, 20 mM Mes<br>50 µM potassium sulfate (2 µCi/mL)<br>2 mM magnesium gluconate<br>**50 mM potassium chloride**<br>pH 5.0 | n = 4 |
| | 20 mM Hepes, 20 mM Mes<br>2 mM magnesium gluconate<br>**50 mM potassium gluconate**<br>pH 7.5 | 20 mM Hepes, 20 mM Mes<br>50 µM potassium sulfate (2 µCi/mL)<br>2 mM magnesium gluconate<br>**50 mM potassium gluconate**<br>pH 5.0 | |
| 1E | 20 mM Hepes, 20 mM Mes<br>50 mM potassium chloride<br>2 mM magnesium gluconate<br>pH 7.5 | 20 mM Hepes, 20 mM Mes<br>50 µM potassium sulfate (2 µCi/mL)<br>45 mM potassium gluconate<br>2 mM magnesium gluconate<br>5 mM **tested anions (sodium salt)**<br>pH 5.0 | n = 4 |
| 1F | 20 mM Hepes, 20 mM Mes<br>42 mM potassium gluconate<br>8 mM potassium chloride<br>2 mM magnesium gluconate<br>pH 7.5 | 20 mM Hepes, 20 mM Mes<br>50 µM potassium sulfate (2 µCi/mL)<br>50 mM potassium gluconate<br>2 mM magnesium gluconate<br>pH 5.0<br><br><u>21-fold counterflow buffer:</u><br>20 mM Hepes, 20 mM Mes<br>50 mM sodium gluconate<br>2 mM magnesium gluconate<br>pH 5.0<br>2.1 mM CCCP<br>**105 mM tested anions (sodium salt)** | n = 3 |
| S2B | 20 mM Hepes, 20 mM Mes (NaOH)<br>**50 mM potassium chloride**<br>2 mM magnesium gluconate<br>pH 7.5 | 20 mM Hepes, 20 mM Mes (NaOH)<br>50 µM potassium sulfate (2 µCi/mL)<br>**50 mM potassium gluconate**<br>2 mM magnesium gluconate<br>100 nM valinomycin<br>pH 5.0 | n = 4 |
| | 20 mM Hepes, 20 mM Mes (NaOH)<br>**5 mM potassium chloride**<br>45 mM sodium chloride<br>2 mM magnesium gluconate<br>pH 7.5 | 20 mM Hepes, 20 mM Mes (NaOH)<br>50 µM potassium sulfate (2 µCi/mL)<br>**5 mM potassium gluconate**<br>45 mM sodium gluconate<br>2 mM magnesium gluconate<br>100 nM valinomycin<br>pH 5.0 | |
| | | 20 mM Hepes, 20 mM Mes (NaOH)<br>50 µM potassium sulfate (2 µCi/mL)<br>**50 mM potassium gluconate**<br>2 mM magnesium gluconate<br>pH 5.0 | |
| | | 20 mM Hepes, 20 mM Mes (NaOH)<br>50 µM potassium sulfate (2 µCi/mL)<br>**5 mM potassium gluconate**<br>45 mM sodium gluconate<br>2 mM magnesium gluconate<br>pH 5.0 | |

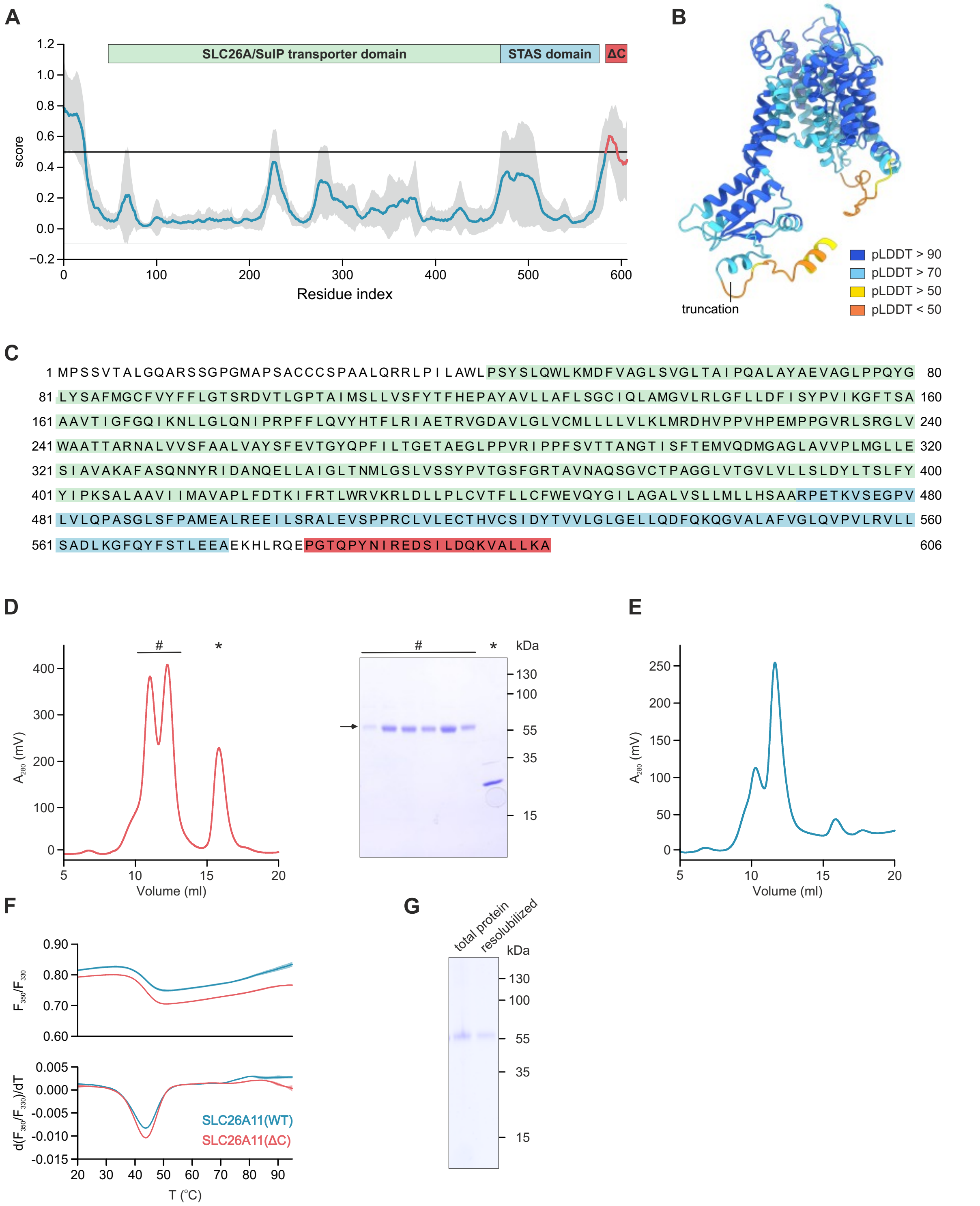


**Supplementary Figure 1: C-terminal truncation of SLC26A11. (A)** Combined intrinsically disordered region prediction for SLC26A11 based on Alphafold 2 pLDDT, IUPRED3, PONDR-VL-XT, PONDR-VL3, PONDR-VSL2 and Disopred3. **(B)** Alphafold model of SLC26A11 with color code showing the pLDDT and position of the C-terminal truncation as indicated. **(C)** SLC26A11 amino acid sequence showing the truncated C-terminus in red. **(D)** Size exclusion of SLC26A11(ΔC) on a Superdex 200 increase 10/300 column and SDS-PAGE of elution fractions showing the monomeric and dimeric species of SLC26A11(ΔC) (#) and the cleaved C-terminal GFP (*). **(E)** Size exclusion of SLC26A11(WT) on a Superdex 200 increase 10/300. **(F)**Thermal unfolding of SLC26A11(WT) and SLC26A11(ΔC). **(G)** SDS-PAGE of SoyPC-SLC26A11(ΔC) proteoliposomes with total protein and DDM resolubilized material recovered from supernatant after high speed ultracentrifugation indicating functional reconstitution of SLC26A11(ΔC).

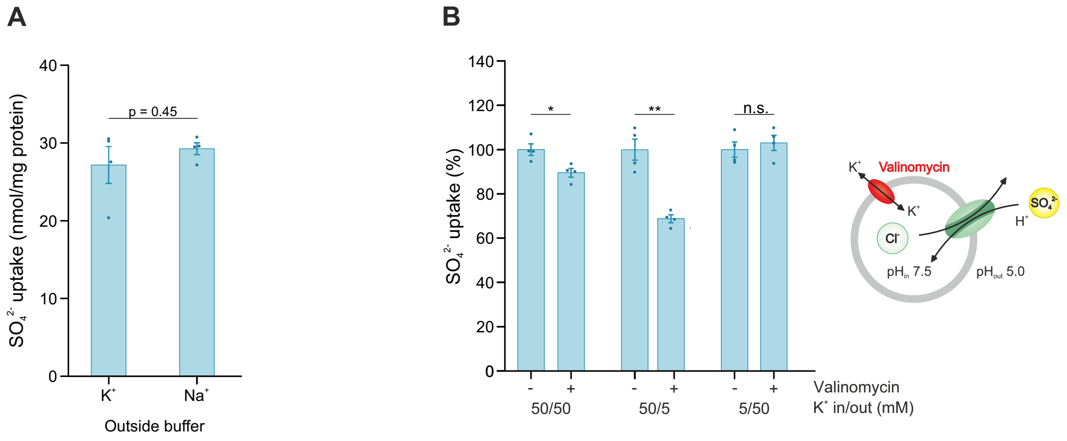


**Supplementary Figure 2: Sulfate transport of SLC26A11$^{\Delta C}$ (A)** $Na^+$-dependence of sulfate transport in proteoliposomes loaded with 50 mM $K^+$ and diluted in buffer containing 50 mM $K^+$ or $Na^+$ as indicated. **(B)** Sulfate transport in presence of different membrane potentials generated by the addition of the $K^+$-ionophore valinomycin and $K^+$-gradients as indicated below the bars. A Two-tailed Student's t test was performed (** $p < 0.01$, * $p < 0.05$, p values are shown in the **Source Data file**). For all experiments, the individual datapoints as well as the mean ± SEM ($n \geq 3$) are shown. Assay buffers and precise number of replicates are detailed in **Supplementary Table 1**. Bar graphs represent sulfate accumulation levels reached after 16 min of transport.

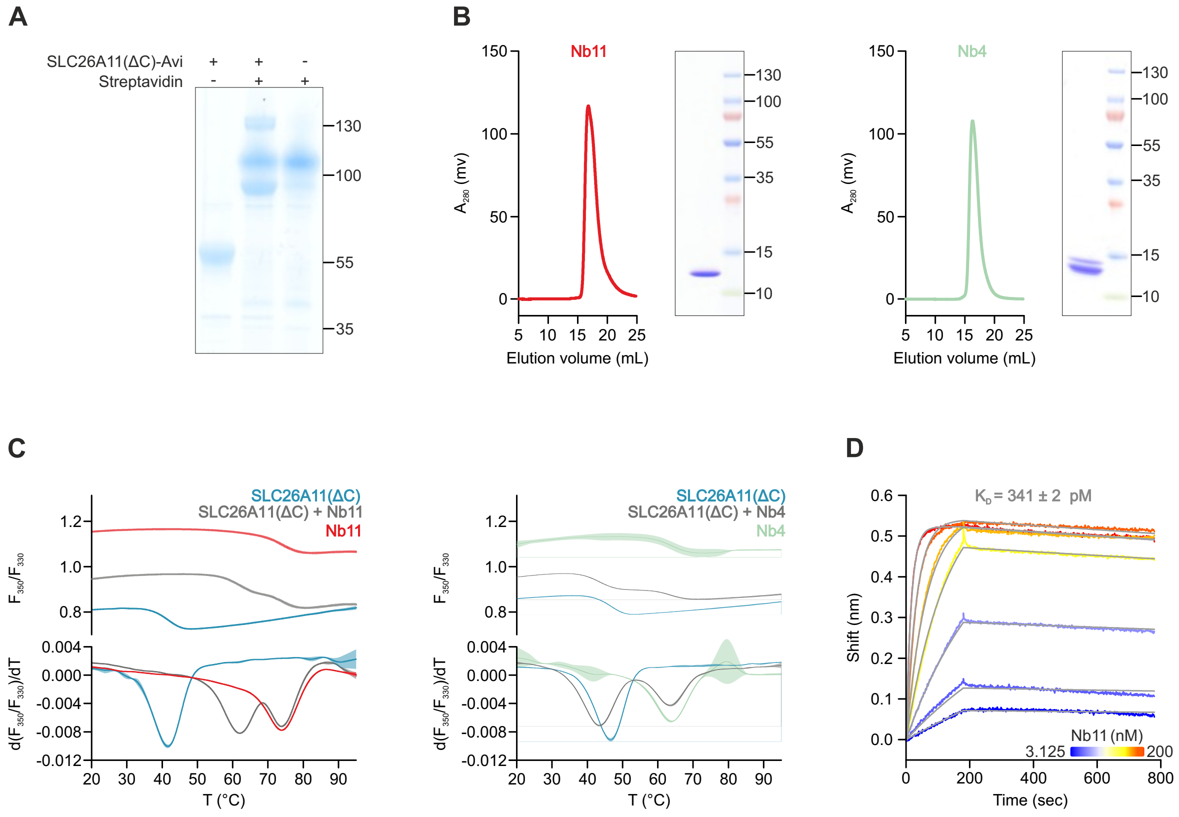


**Supplementary Figure 3: Nanobody selection, purification and characterization. (A)** Mobility shift in SDS-PAGE of enzymatically biotinylated SLC26A11(ΔC)-Avi in presence of Streptavidin. **(B)** Size exclusion chromatogram (Sepax SRT-10C SEC-300 column) and SDS-PAGE of purified Nb11 and Nb4. **(C)** Thermal unfolding of SLC26A11(ΔC) in presence of Nb11 or Nb4. **(D)** Bio-layer interferometry based affinity determination of Nb11 binding to immobilized SLC26A11(ΔC)-Avi.

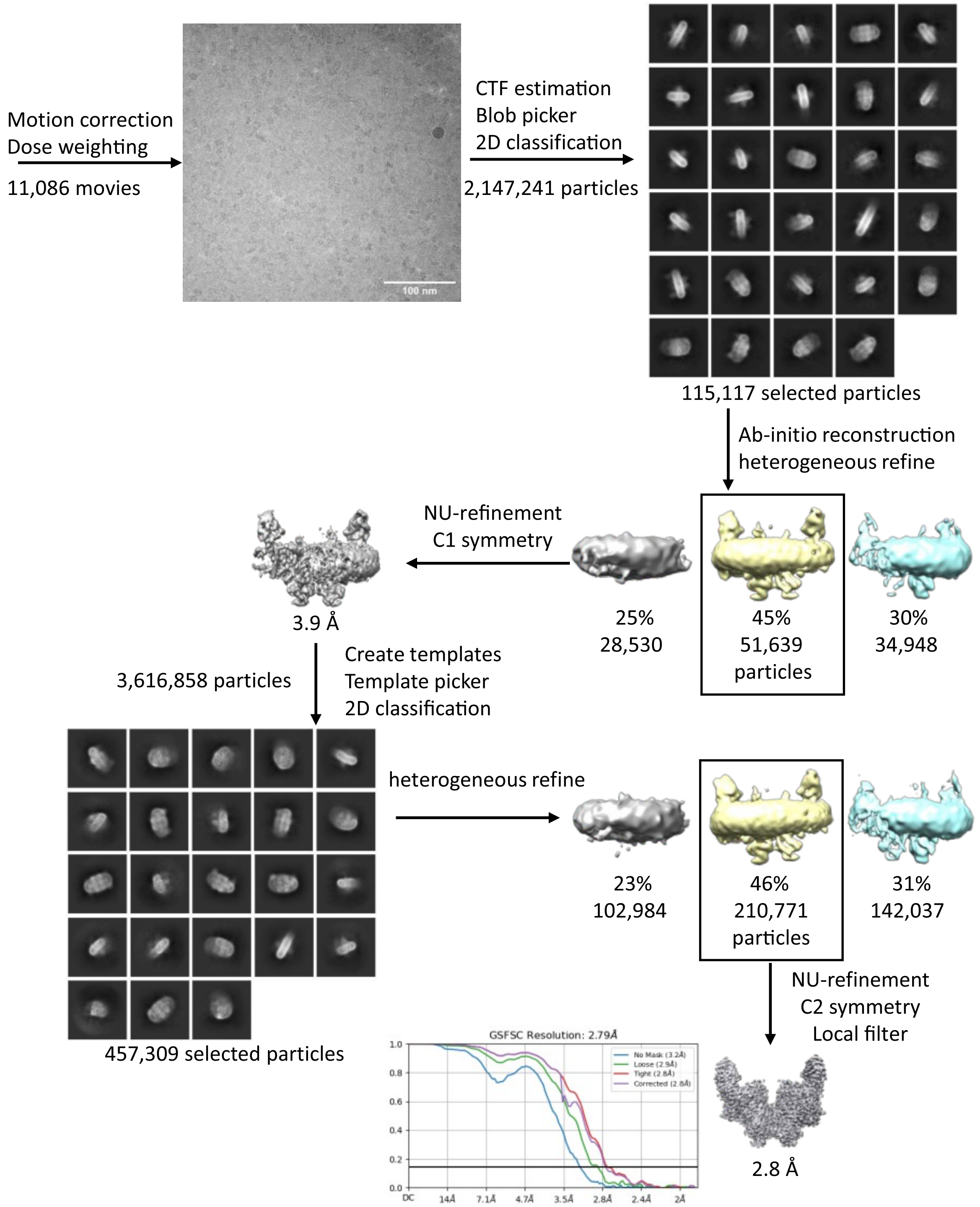


**Supplementary Figure 4: Cryo EM workflow of SLC26A11 with Nb4.**

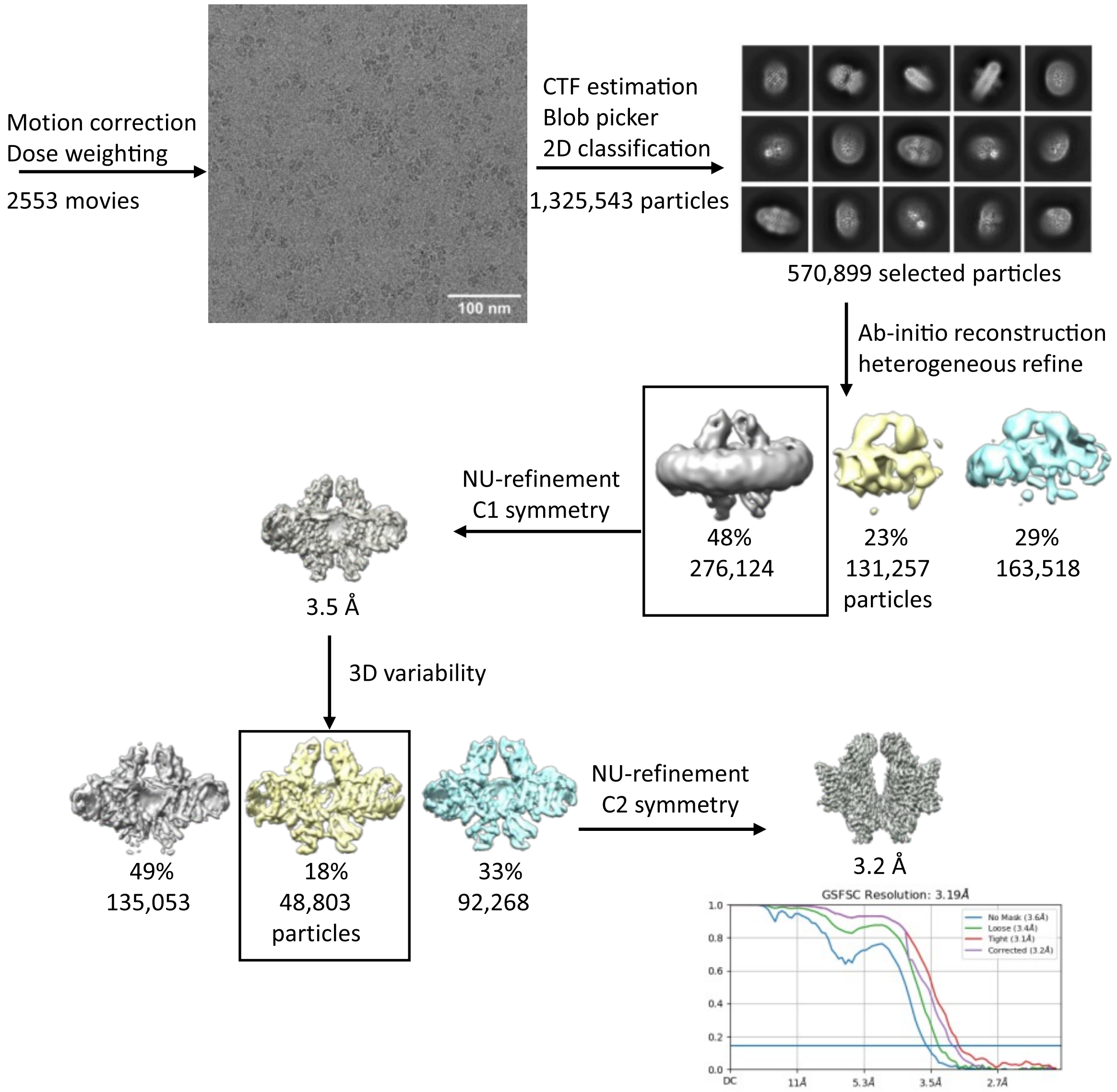


**Supplementary Figure 5: Cryo EM workflow of SLC26A11 with Nb11.**

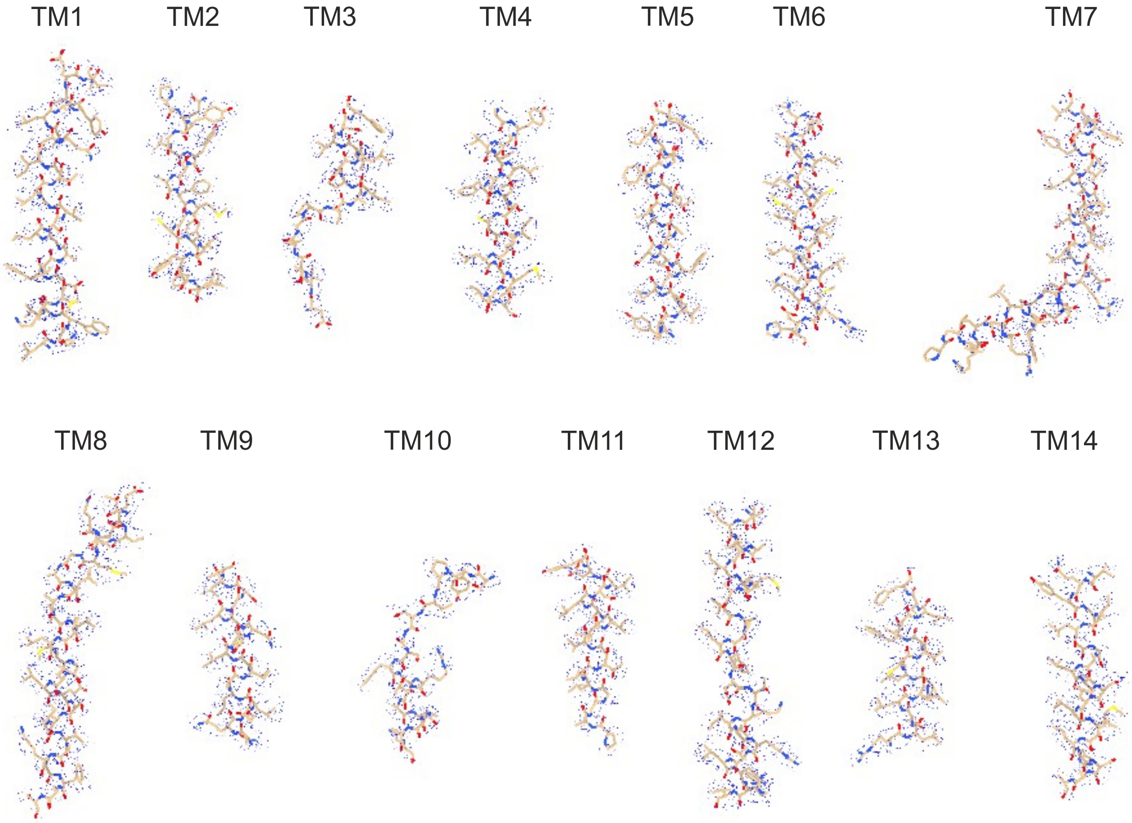


**Supplementary Figure 6: Representative densities of SLC26A11-Nb11.** Transmembrane segments shown as sticks with corresponding density (blue mesh).

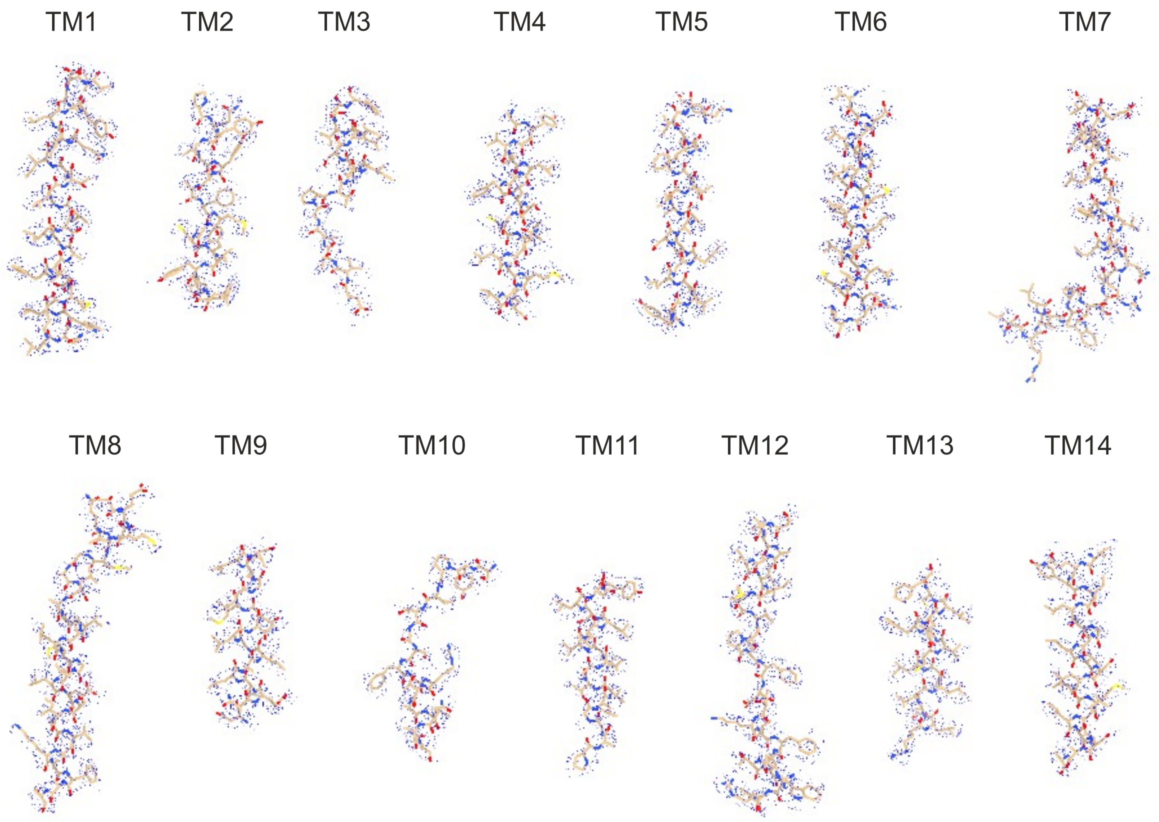


**Supplementary Figure 7: Representative densities of SLC26A11-Nb4.** Transmembrane segments shown as sticks with corresponding density (blue mesh).

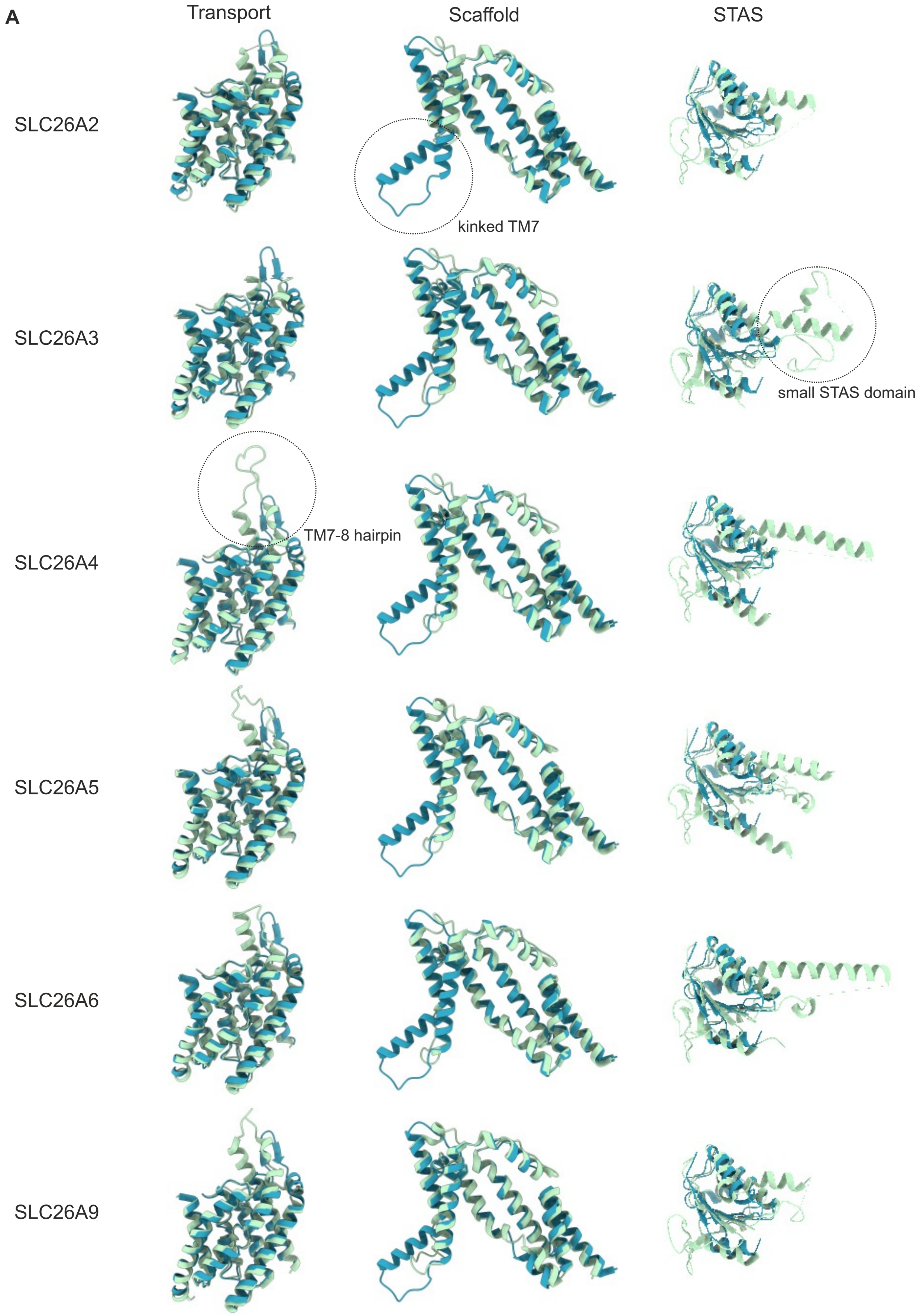
A
Transport
Scaffold
STAS
SLC26A2
kinked TM7
SLC26A3
small STAS domain
SLC26A4
TM7-8 hairpin
SLC26A5
SLC26A6
SLC26A9

B

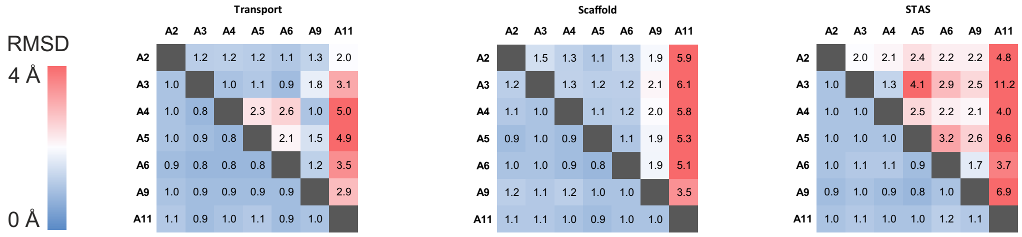


**Supplementary Figure 8: The three unique structural motifs of SLC26A11. (A)** Structural alignment based on the matchmaker command in UCSF ChimeraX of the individual domains of SLC26A11 (Transport, Scaffold and STAS, blue) with domains from all other human SLC26 isoforms (green) with available experimental high resolution structure. Unique structural features of SLC26A11 are highlighted by black circles. **(B)** RMSD from structural alignment of individual domains of human SLC26 isoforms in ChimeraX. Upper right shows results from all atom pairs and lower left from pruned atom pairs.

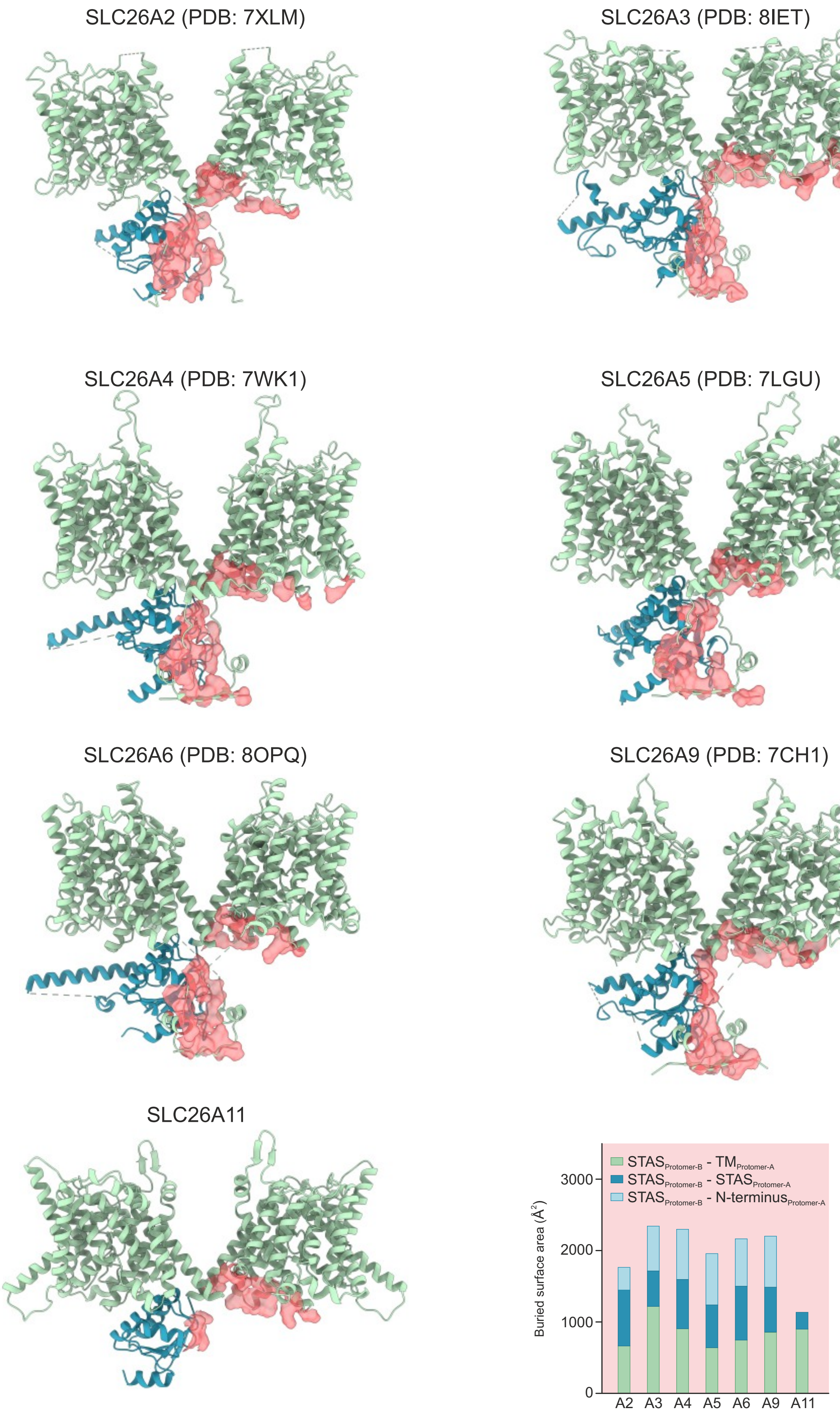


**Supplementary Figure 9: STAS mediated dimer interface of human SLC26 transporters.** Structures of human SLC26 isoforms with available experimental high-resolution structure with transmembrane domains in green and STAS domain in blue and the second STAS domain left out for clarity. Interfaces are indicated as red surface. Total interface size and contribution of transmembrane domain, STAS domain and N-terminus to the interface were quantified using the interfaces command in UCSF ChimeraX.

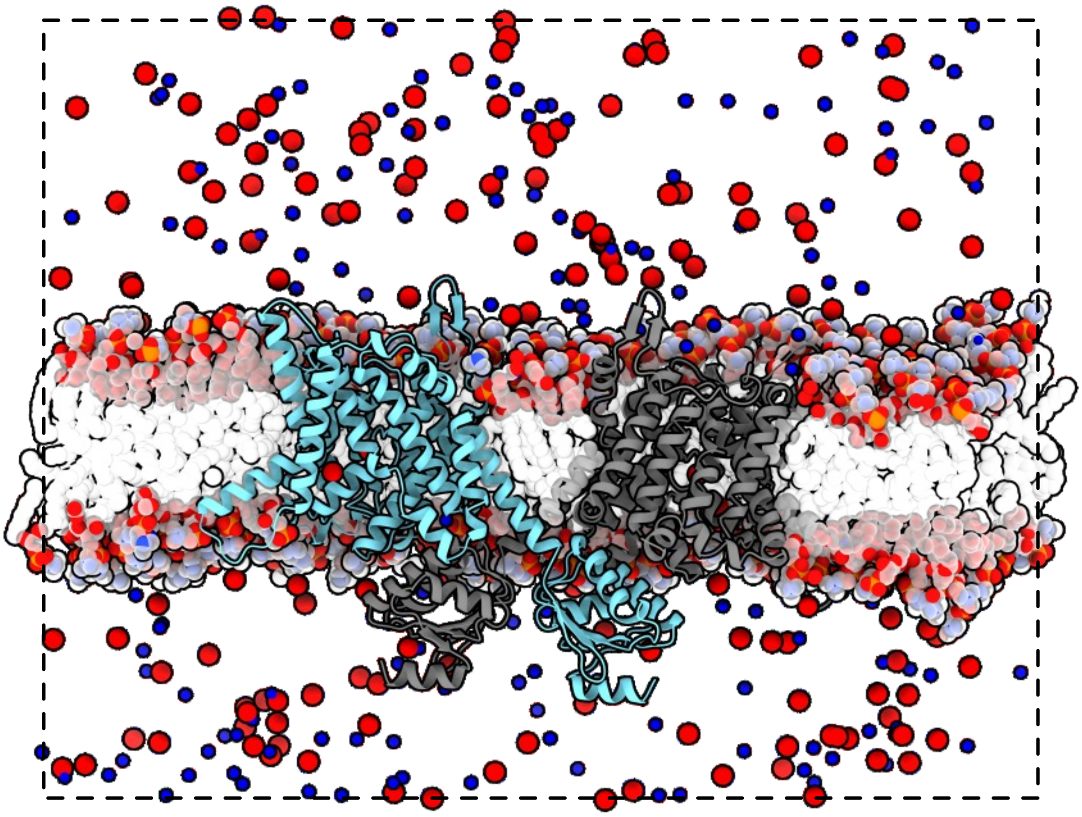

**Supplementary Figure 10: SLC26A11 simulation system.** Simulation system of SLC26A11 embedded in a POPC membrane, fully solvated (not shown) and ionized in ~200 mM $Na^+$ (blue)/$Cl^-$ (red).

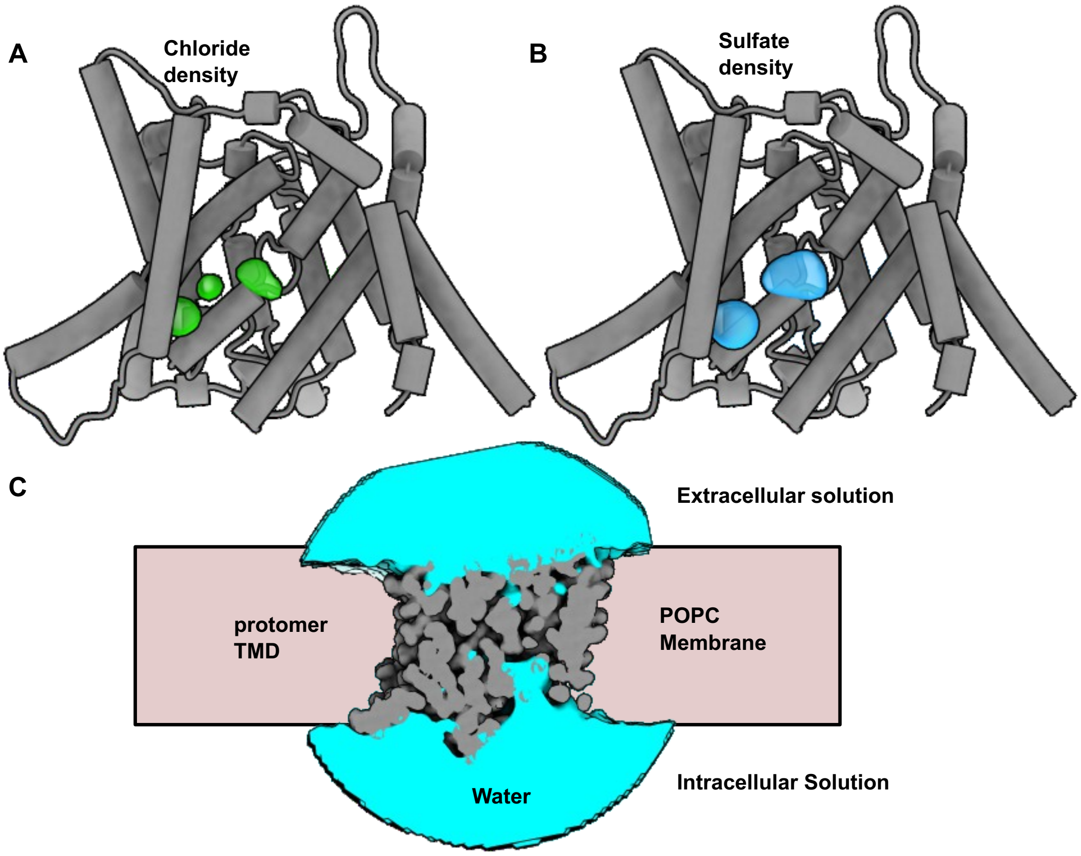


**Supplementary Figure 11: Chloride and sulfate densities.** Total averaged chloride **(A)** and sulfate **(B)** densities calculated from equilibrium MD simulations. Densities are shown at an iso-density threshold of 0.1 AMU/Å3. Densities are shown against the transmembrane domain of a single protomer. **(C)** Average water density around a TMD protomer.

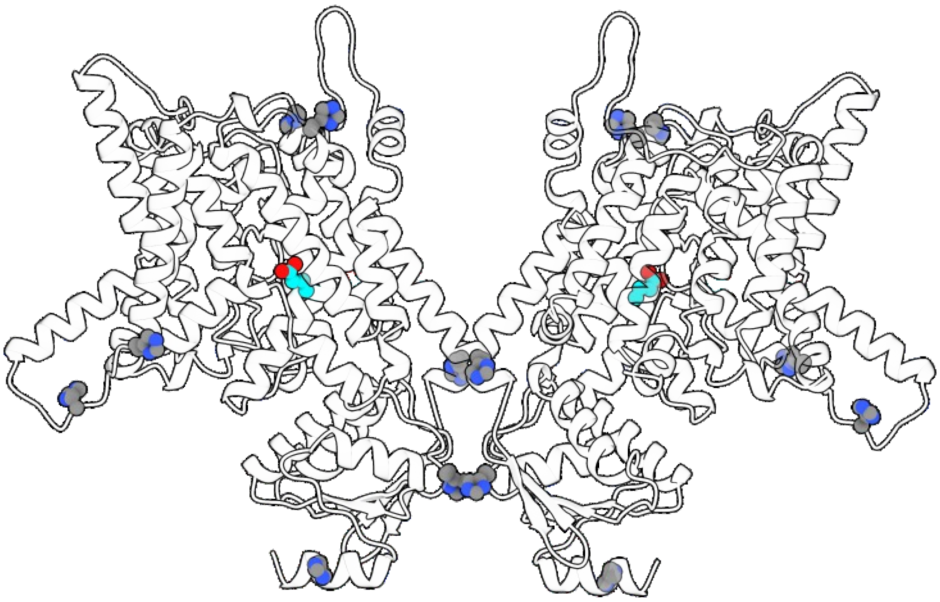

**Supplementary Figure 12: Dimeric structure of SLC26A11 in cartoon representation.** Location of histidine side chains exclusively on the intra and extra cellular sides of the protein is highlighted by grey sphere representation. The location of glutamate-320 is illustrated in cyan.

**A**

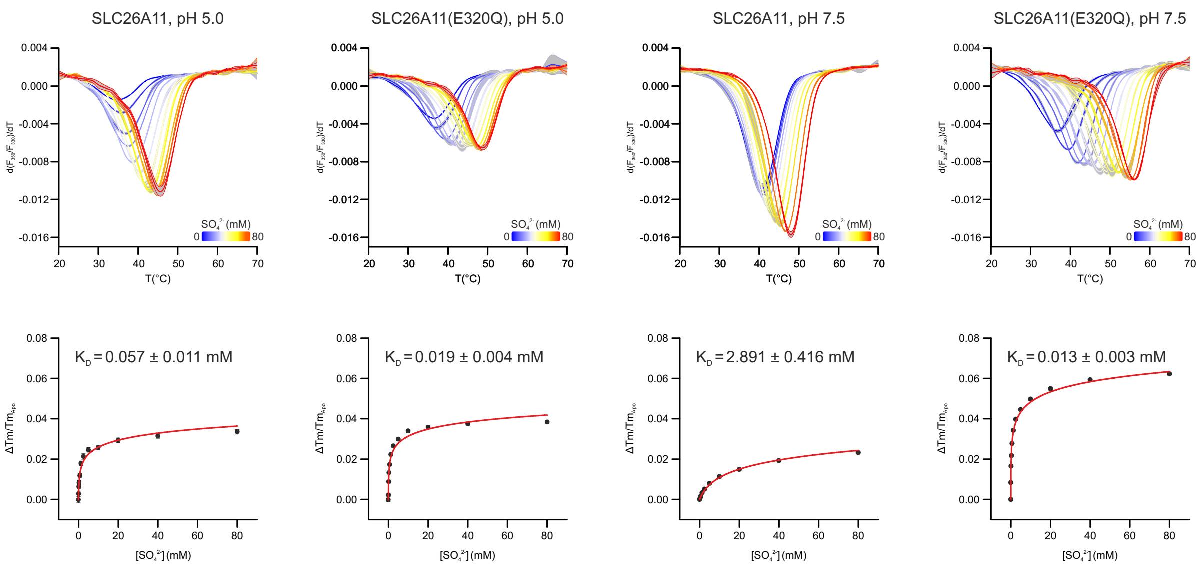


**B**

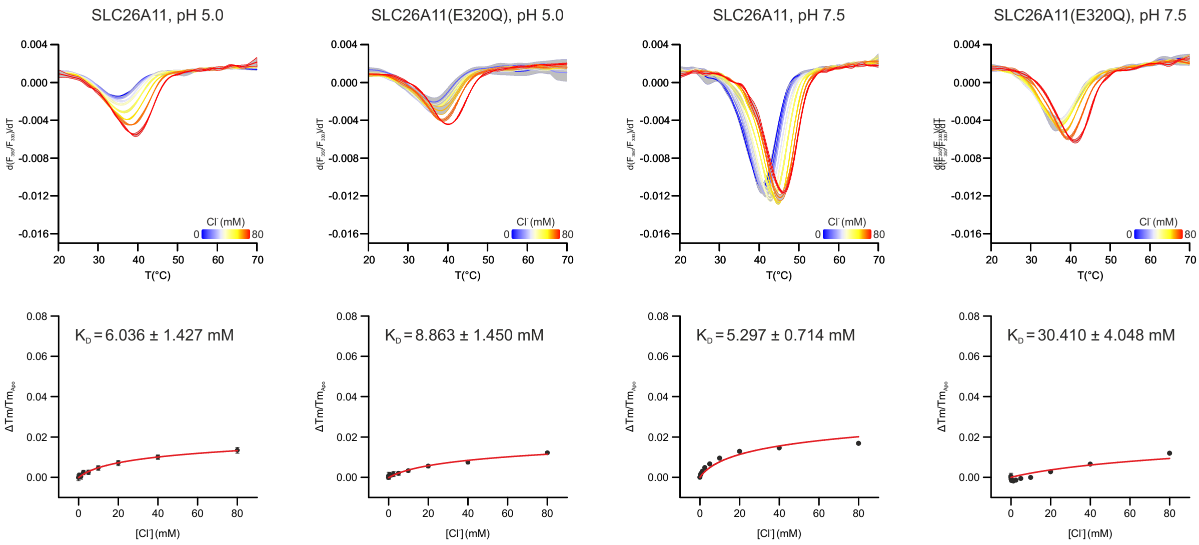


**Supplementary Figure 13: pH dependent substrate binding. (A)** Thermal unfolding of SLC26A11(ΔC) and SLC26A11(ΔC, E320Q) at pH 5.0 or pH 7.5 in presence of increasing concentrations of sulfate (blue = 0 mM sulfate, red = 80 mM sulfate) and derived dissociation constants ($K_D$). **(B)** Thermal unfolding of SLC26A11(ΔC) and SLC26A11(ΔC, E320Q) at pH 5.0 or pH 7.5 in presence of increasing concentrations of chloride (blue = 0 mM chloride, red = 80 mM chloride) and derived dissociation constants ($K_D$). Shown are mean and standard deviation (n = 3) of the first derivative of $F_{350}/F_{330}$. The melting temperature Tm is reported by the local minimum of $d(F_{350}/F_{330})/dT$.

**A**

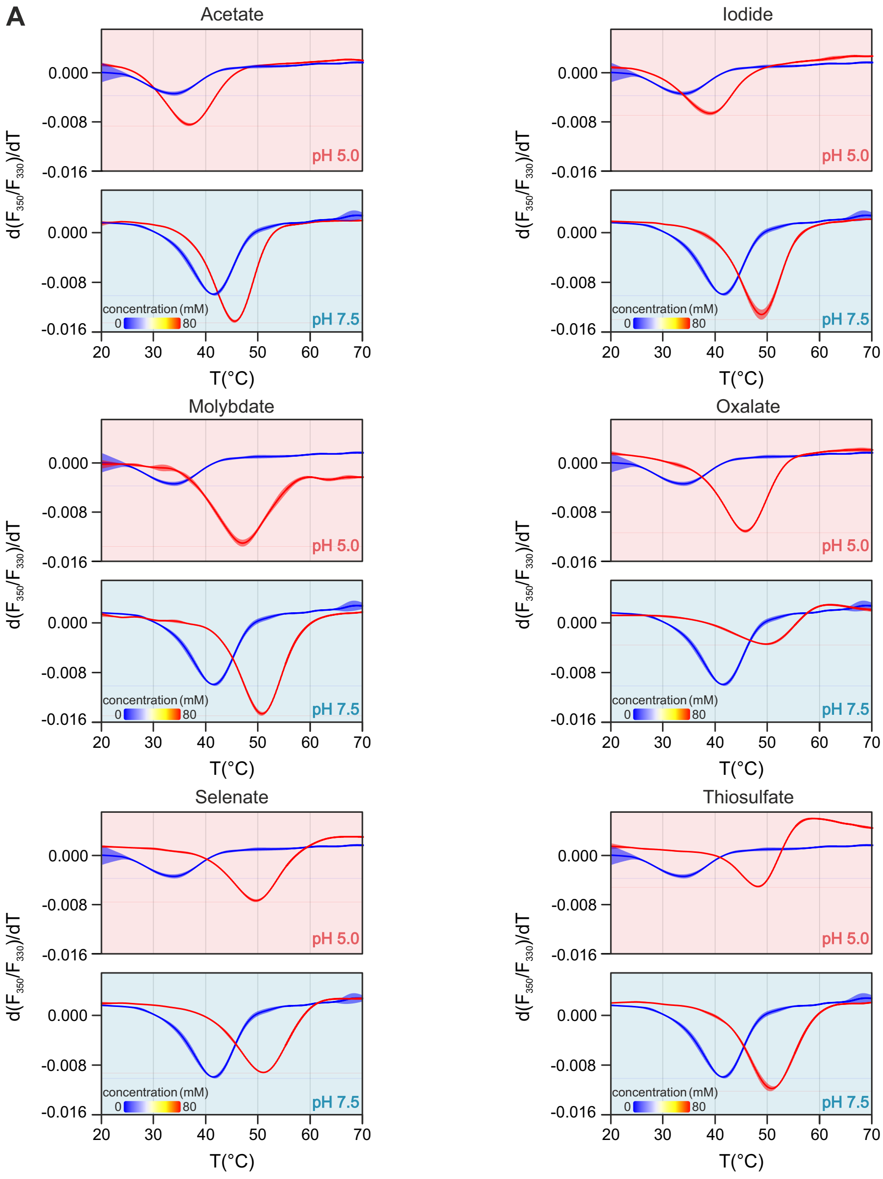


**B**

| | pH 5.0 $K_D$ (mM) | pH 7.5 $K_D$ (mM) |
|---|---|---|
| Thiosulfate | 0.009 ± 0.003 | 1.189 ± 0.365 |
| Selenate | 0.004 ± 0.001 | 1.121 ± 0.248 |
| Oxalate | 0.040 ± 0.011 | 2.110 ± 0.594 |
| Molybdate | 0.017 ± 0.005 | 1.241 ± 0.302 |
| Iodide | 2.950 ± 0.567 | 3.016 ± 0.640 |
| Acetate | 12.426 ± 1.393 | 15.522 ± 1.690 |

**C**

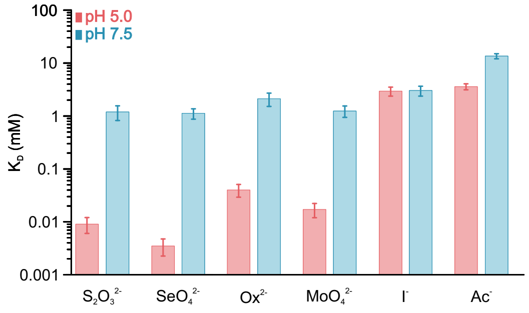


**Supplementary Figure 14: pH dependent anion binding. (A)** Thermal unfolding of SLC26A11(ΔC) in presence of 80 mM anion added as sodium salt at pH 5.0 or pH 7.5, respectively, with mean and S.D. (n = 3). **(B)** pH dependent dissociation constants ($K_D$) of tested anions calculated from data shown in panel A. **(C)** Bar graph of data from panel B.

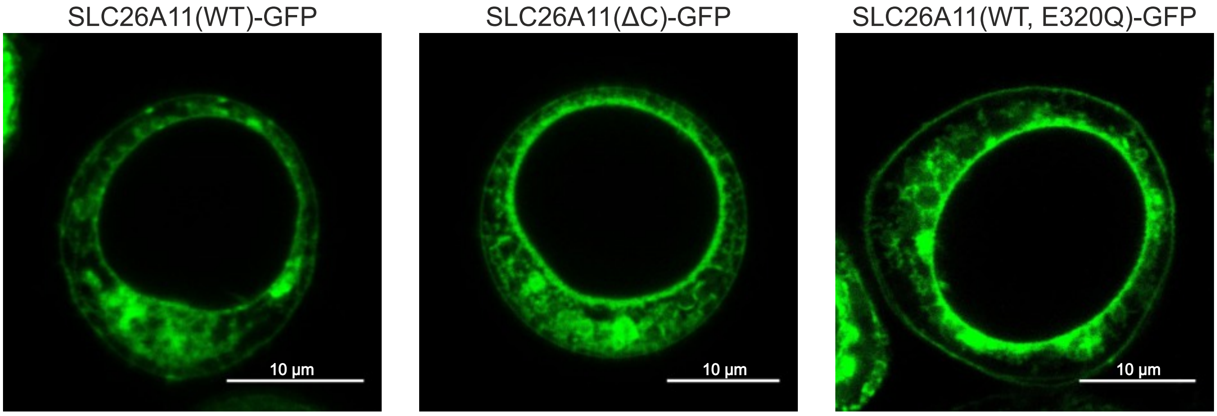


**Supplementary Figure 15: Expression of SLC26A11-GFP constructs in Sf9 cells.** Confocal microscopy of Sf9 cells transfected with either SLC26A11(WT), SLC26A11(ΔC) or SLC26A11(WT,E320Q) as indicated. All SLC26A11 variants show fractional localization to the plasma membrane with the majority of proteins localized to intracellular compartments.

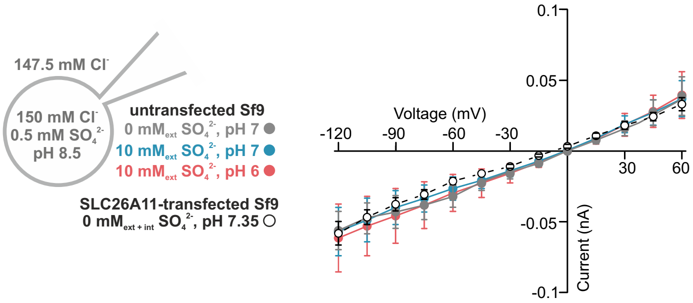


**Supplementary Figure 16:** Current-voltage relationships from untransfected *Sf*9 cells in three different bath solutions as indicated and transfected *Sf*9 cells in absence of internal sulfate. Shown are means and standard errors.

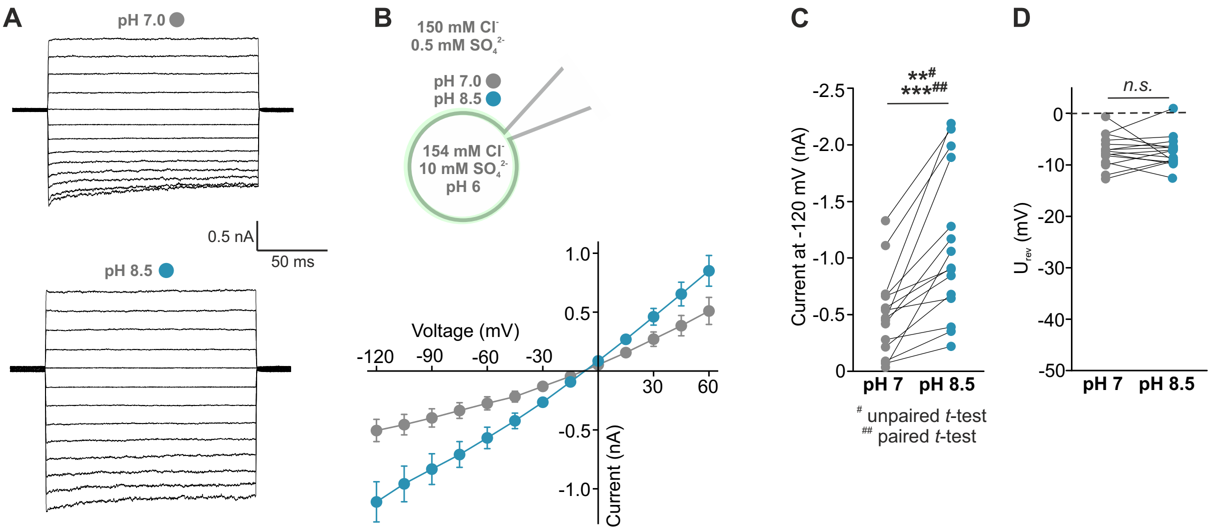


**Supplementary Figure 17: Alkanisation stimulates channel currents during sulfate export.** (**A**) Representative current recordings from *Sf*9 cells expressing SLC26A11(WT)-eGFP with buffer conditions as shown in the inset. (**B**) Current-voltage relationships from transfected *Sf*9 cells in two different bath solutions as indicated (grey: 140 mM $Cl^-$, pH 7; blue: 140 mM $Cl^-$, pH 8.5, n=15/15). Shown are means and standard errors. **(C)** Statistical analysis of current amplitudes at -120 mV for all experiments. An unpaired *t*-test show that means of current amplitudes are significantly different (p=0.004) and pairwise comparison show, that current stimulation by alkanisation was observed in all cases (p<0.001), whereas reversal potentials (**D**) remained constant (p=0.898).

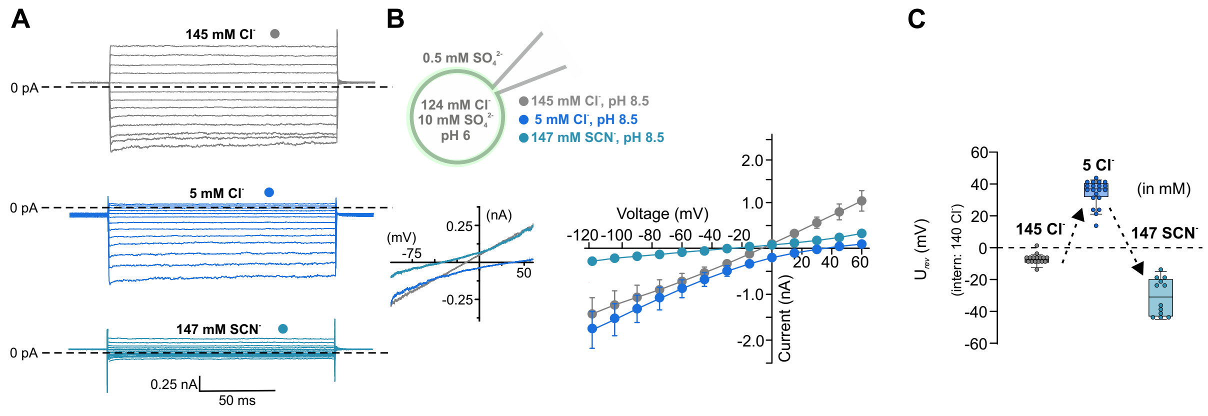


**Supplementary Figure 18: SLC26A11 channel is anion-selective. (A)** Representative current recordings from *Sf*9 cells expressing SLC26A11(WT)-eGFP with buffer conditions as shown in the inset. **(B)** Current-voltage relationships from transfected *Sf*9 cells in three different bath solutions as indicated (grey: 145 mM $Cl^-$, pH 8.5; blue: 5 mM $Cl^-$, pH 8.5; cyan: 147 mM $SCN^-$, pH $8.5_{ext}$:$6.0_{int}$). Shown are means and standard errors. The inset in B shows current recordings from voltage ramps from a representative experiment. **(C)** Reversal potentials ($U_{rev}$) from experiments shown in panel A and B for WT (145 mM $Cl^-$, 5 mM $Cl^-$, 147 mM $SCN^-$: n=16/20/12).

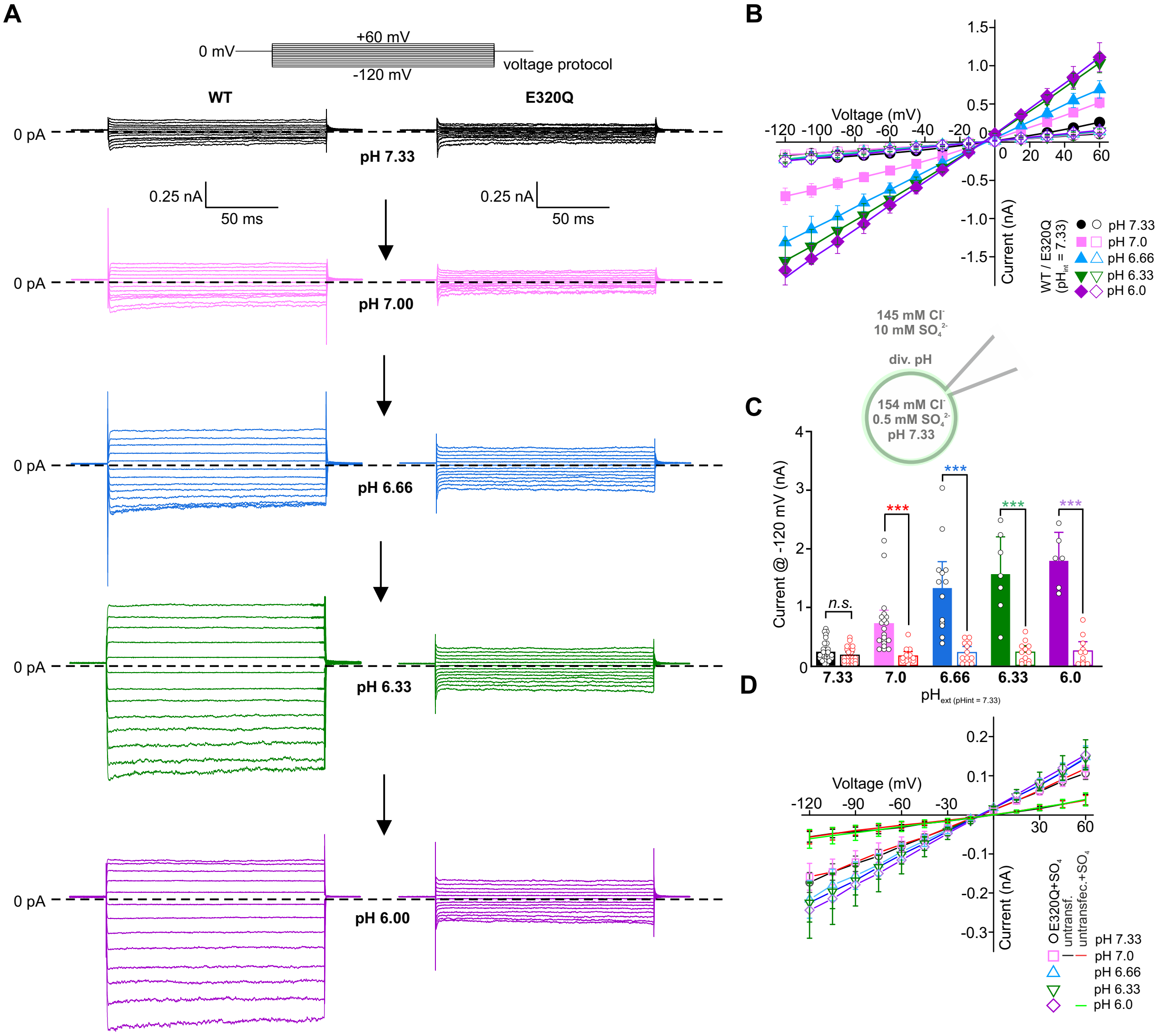


**Supplementary Figure 19: Stimulation of SLC26A11 channel by external pH is abolished in E320Q mutant.** (**A**) Representative current recordings from *Sf*9 cells expressing SLC26A11(WT)-eGFP (left) and SLC26A11(E320Q)-eGFP with buffer conditions as indicated in the inset. (**B**) Current-voltage relationships from transfected *Sf*9 cells (WT: closed symbols; E320Q: open symbols) in five different bath solutions as indicated (black: pH 7.33; magenta, pH 7.0; blue: pH 6.6, green: pH 6.33, violet: pH 6.0). **(C)** Statistical analyses of current amplitudes at -120 mV for all tested external pH values (*2-way* ANOVA with Holm-Sidak *posthoc* testing, $p<0.001$). (**D**) Current-voltage relationships of background currents from *Sf9* cells (lines, no symbols) and E320Q-mediated residual currents (lines, open symbols) for different ionic conditions. Shown are means and standard errors.

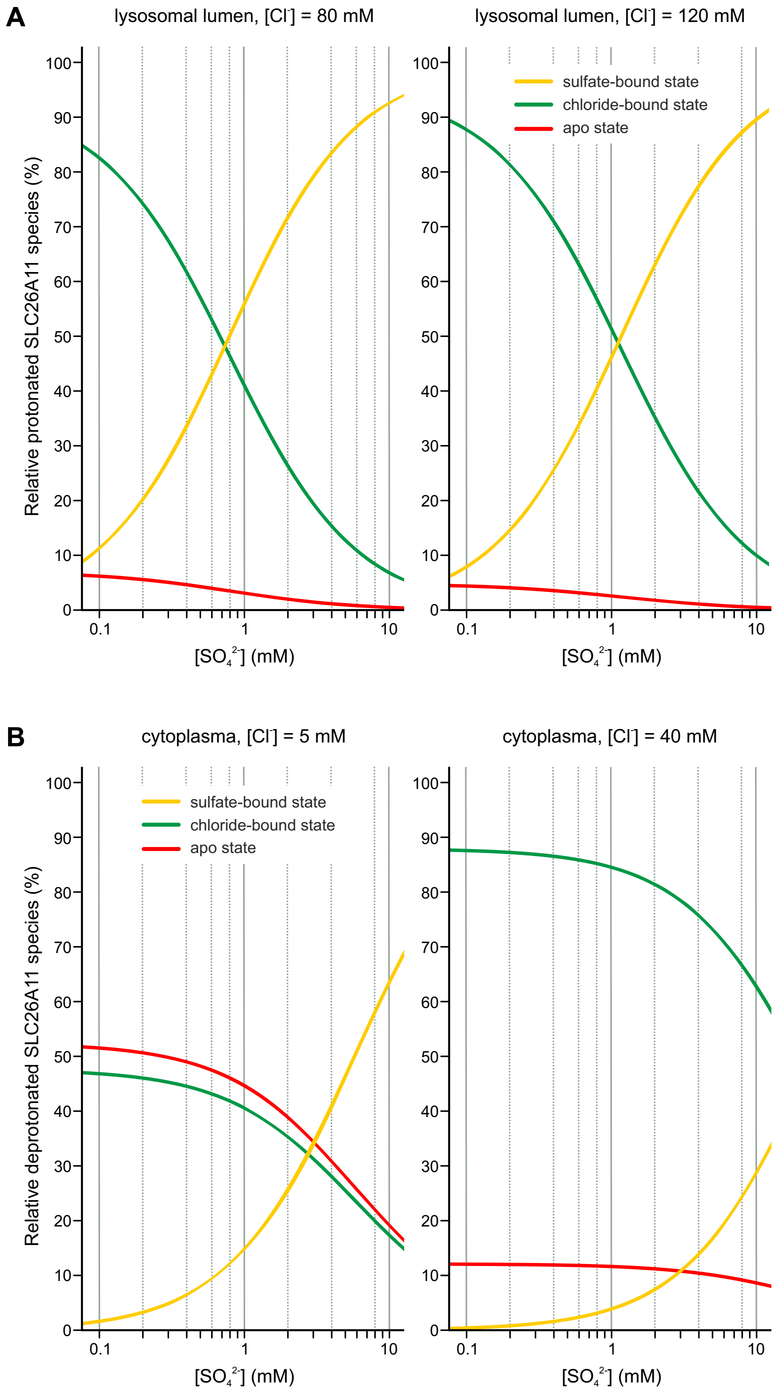


**Supplementary Figure 20: Modeling of competitive binding of sulfate and chloride to SLC26A11.** (**A**) Competitive binding of sulfate and chloride to the SLC26A11 substrate binding site exposed to the lumen of the lysosome and with Glu-320 in the protonated form. The fraction of each SLC26A11 species (sulfate-bound, chloride-bound, or apo) is depicted as a function of the lysosomal sulfate concentration. The model is based on the experimentally determined dissociation constants (sulfate: $K_D$ 57 µM; chloride: $K_D$ 6.0 mM) and luminal chloride concentrations of 80 mM (left panel) and 120 mM (right panel). (**B**) Competitive binding of sulfate and chloride to the SLC26A11 substrate binding site exposed to the cytoplasm and with Glu-320 in the deprotonated form. The fraction of each SLC26A11 species (sulfate-bound, chloride-bound, or apo) is depicted as a function of the lysosomal sulfate concentration. The model is based on the experimentally determined dissociation constants (sulfate: $K_D$ 2.9 mM; chloride: $K_D$ 5.3 mM) and cytoplasmic chloride concentrations of 5 mM (left panel) and 40 mM (right panel). Models were generated using the simulation applet from Pääkkonen *et al.*, 2022 (https://doi.org/10.1021/acsomega.2c00560).
